# Proteoform-level deconvolution reveals a broader spectrum of ibrutinib off-targets

**DOI:** 10.1101/2023.11.14.566837

**Authors:** Isabelle Leo, Elena Kunold, Audrey Anastasia, Marianna Tampere, Jürgen Eirich, Rozbeh Jafari

**Affiliations:** Clinical Proteomics Mass Spectrometry, Department of Oncology-Pathology, Karolinska Institutet, Science for Life Laboratory, Solna, Sweden; Department of Medical Oncology, University Medical Center Groningen, Groningen, Netherlands; Precision Cancer Medicine, Department of Oncology-Pathology, Karolinska Institutet, Science for Life Laboratory, Solna, Sweden; Institute of Plant Biology and Biotechnology, University of Munster, Munster, Germany

## Abstract

Over the last decade, proteome-wide mapping of drug interactions has revealed that most targeted drugs bind to not only their intended targets, but additional proteins as well. However, the majority of these studies have focused on analyzing proteins as encoded by their genes, thus neglecting the fact that most proteins exist as dynamic populations of multiple proteoforms. Here, we addressed this problem by combining the use of thermal proteome profiling (TPP), a powerful method for proteome analysis, with proteoform detection to refine the target landscape of an approved drug, ibrutinib. We revealed that, in addition to known targets, ibrutinib exhibits an intricate network of interactions involving multiple different proteoforms. Notably, we discovered affinity for specific proteoforms that link ibrutinib to mechanisms in immunomodulation and cellular processes like Golgi trafficking, endosomal trafficking, and glycosylation. These insights provide a framework for interpreting clinically observed off-target and adverse events. More generally, our findings highlight the importance of proteoform-level deconvolution in understanding drug interactions and their functional impacts, and offer a critical perspective for drug mechanism studies and potential applications in precision medicine.

## Introduction

Proteins are responsible for execution of cellular processes and act as critical mediators of phenotypic traits. Although the human genome encodes approximately 20,000 unique protein sequences, a multitude of processes including alternative splicing, post-translational modifications (PTMs), and proteolytic cleavage serve to significantly expand this diversity, generating proteoforms in numbers several orders of magnitude higher (*1*). These proteoforms open the door to much wider range of function and evolution of adaptive phenotypes under selective pressure(*1*). Additionally, many diseases, such as cancer, have been linked to changes that result in alterations to proteoforms, such as decay of aberrant transcripts(*2*), splicing dynamics(*3*), translation(*4*), and post-translational modifications(*5*). With highly variable proteomes that are under continuous selective pressure, cancers demonstrate numerous examples where protein complex composition(*6–8*) and proteoform drift(*9–12*) play important roles in biology and therapeutic response. Therefore, targeting cancer-associated proteoforms has emerged as an area of intense clinical interest, notably in pediatric cancers(*13*), prostate cancer(*14*), melanoma(*15*, *16*), and breast cancer(*17*). This highlights the importance of identifying and characterizing different proteoforms, as well as developing drugs that target them. These efforts have the potential to significantly contribute to advancing precision medicine and delivering on the promise of personalized therapy.

In this context, development of targeted therapies for treatment of cancer has been one of the major breakthroughs in the field. These advances have been fueled by our improved ability to connect a disease phenotype to a specific causative protein, and then translate these insights into a drug that targets that protein to correct its function and restore the phenotype. Although the focus of targeted drug discovery is on development of drugs that affect only a single target, global proteomic mapping of drug-proteome interactions have revealed that many approved targeted therapies affect more than a single target protein (*18*, *19*). These off-target effects can often help rationalize clinical observations both with respect to efficacy and toxicity. However, the full understanding of how drugs bind and affect different proteoforms has been missing, given the difficulty of conducting proteoform-level analysis.

In general, proteoform species are challenging to detect and quantify because they lack a one-to-one relationship with genetic information, making their existence hard to predict and anticipate. Additionally, proteoforms may occur at low levels, or only transiently. Thus, molecular methods for proteoform characterization have to allow for unbiased identification without a need to pre-define or isolate variants, and offer high sensitivity and dynamic range. Currently, both top-down and bottom-up mass spectrometry (MS)-based proteomics have been used to conduct proteoform analysis. For example, top-down proteomics has been employed to assemble human proteoform reference maps(*20*, *21*); however, this method has limitations in depth and throughput(*21*). In contrast, bottom-up proteomics uses digested peptide libraries to achieve the broadest range of identifications. This makes it suitable for global proteome detection approaches(*22*), but at the cost of inferring proteoforms rather than identifying them directly.

To address this issue, many methods for global proteoform inference have been developed, including thermal proteome profiling (TPP)(*23*). TPP is a powerful method for systematic detection and annotation of functional aspects of the proteome that uses a series of temperature treatments to resolve proteins based on their thermal stability(*24*). Due to varying thermal stability of proteoforms, which can arise from different PTMs, alternative splicing or proteolytic processing, and interactions with proteins, DNA, RNA, metabolites or drugs, TPP can represent many proteoform types(*23*). Furthermore, this method can be applied to a wide range of biological systems and used to analyze proteoforms in their native contexts.

Here, we used TPP to describe small molecule drug interactions with proteoforms. We selected to focus on mapping the target landscape of ibrutinib, a clinically used Bruton tyrosine kinase (BTK) inhibitor(*25–27*). Our choice to focus on ibrutinib was motivated by clinical data that point to a complex relationship between on-target binding, efficacy and toxicity. For example, ∼85% of chronic lymphocytic leukemia (CLL) patients treated with ibrutinib develop BTK pathway mutations (*28–30*). However, some of these patients continue to respond to ibrutinib(*31*, *32*), which could be attributed to a range of factors, including possibly inhibition of additional targets. Although previous proteomics studies have identified a wide range of ibrutinib off-targets(*18*, *33*), which could be clinically beneficial and useful for repurposing (*34*, *35*) or harmful to patients(*41*, *42*), many clinical observations remain to be rationalized. Therefore, we hypothesized that proteoform-level analysis would provide a more complete picture of the ibrutinib target landscape and extend our understanding of treatment sensitivity, off-target events, and resistance mechanisms. Our study revealed additional targets for ibrutinib, including proteoforms involved in Golgi trafficking, glycosylation, cell adhesion, and endosomal processing, as well as some that may amplify drug efficacy and enable BTK-independent immunomodulation.

## Results

### Proteoform identification in ibrutinib treated cell lysates

To reduce the risk of ambiguity in distinguishing drug targets from indirect secondary alterations, we performed TPP in cell lysates. We used lysates from two different cells lines, a precursor-B acute lymphoblastic leukemia cell line with a somatic form of BTK, RCH-ACV(*36*) (RRID: CVCL_1851), and the adrenal carcinoma cell line SW-13 also with somatic BTK(*36*) (RRID: CVCL_0542). Lysates were treated with 100uM high dose ibrutinib or equivalent volume of DMSO, and each lysate and treatment condition was prepared in technical replicates for a total of 8 temperature melt curve sets. Each set was thermally denatured in a 10-point temperature curve and pre-fractionated using high-resolution isoelectric focusing(*37*) for in-depth peptide detection by MS proteomics. In total, 175379 unique peptides mapping to only one gene symbol were detected from 11043 gene symbol stratified proteins. After proteoform clustering analysis(*23*) of these peptides across the whole dataset, 16079 proteoforms were identified. These proteoforms were investigated for differential melting using nonparametric analysis of response curves (NPARC)(*38*), considering proteoforms detected in all samples first across the entire dataset and then within each lineage separately (Figure 1A, Supplementary Figure 1A). Together, these analyses identified 2305 thermally impacted proteoforms from 1936 gene symbols at a p-value threshold of 0.05 (Supplementary Table 1), and 251 proteoforms from 230 gene symbols at a Benjamini-Hochberg adjusted p-value threshold (pAdj) of 0.05.

**Figure 1.**
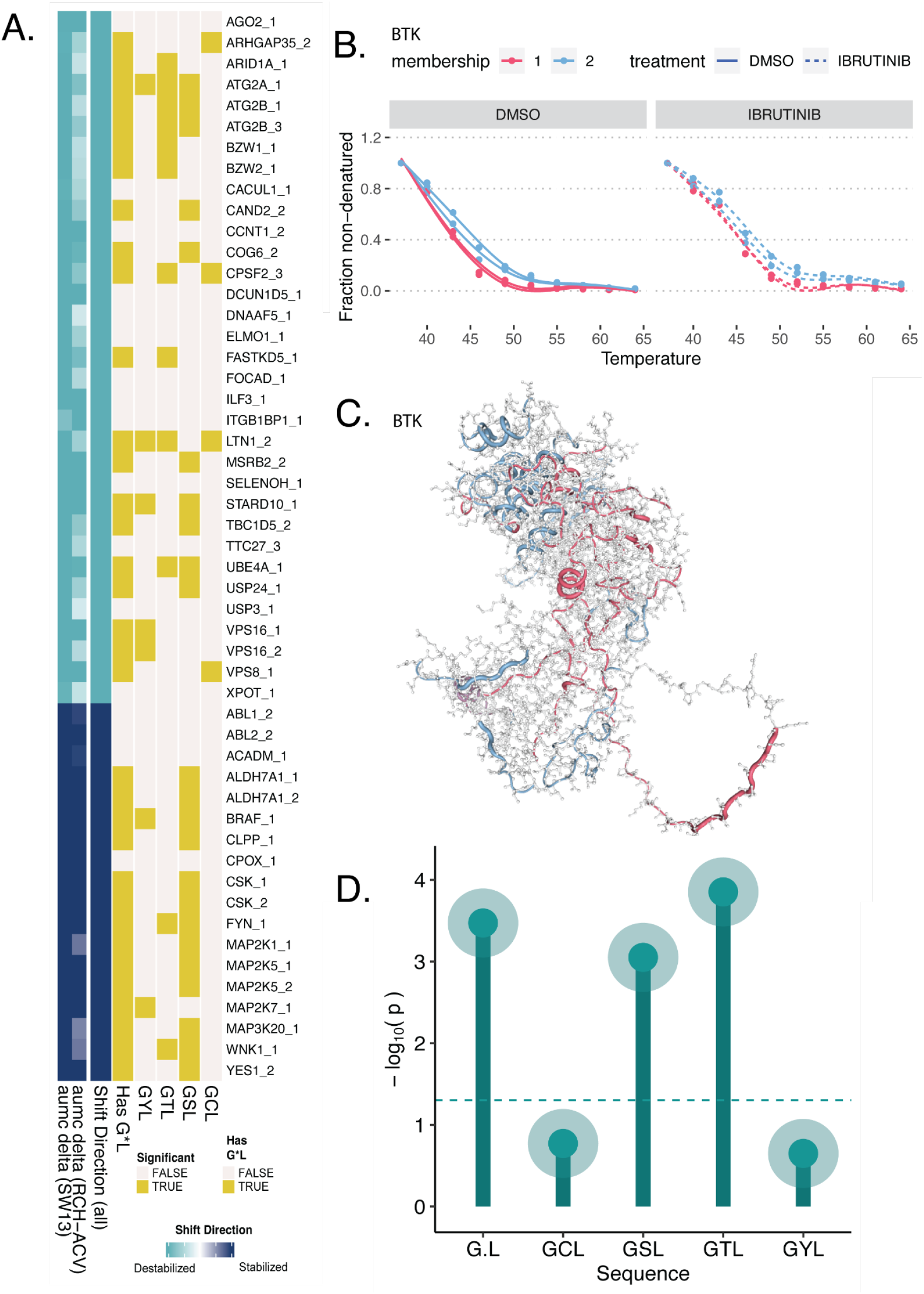
Characteristics of top ibrutinib binding candidates and BTK proteoforms: **A)** The top 51 proteoform results differentially melting in ibrutinib treated versus untreated lysates, as determined by NPARC analysis, with highlighted GCL, GSL, GTL, or GYL sequences. Selected proteoforms meet a Benjamini-Hochberg adjusted p-value threshold of less than 0.0001 for models with detection in all cell lines. **B)** Fraction non-denatured for each BTK proteoform detected in RCH-ACV, colored by proteoform membership assignment demonstrating melting behavior by proteoform and treatment status. **C)** Structural diagram of peptide mappings, generated using the Alphafold structure for BTK generated using the canonical FASTA sequence (sp|Q06187|BTK_HUMAN), showing tube overlays for peptides colored by their proteoform assignments. Regions with multiple matched proteoforms are displayed with blended translucent coloring, and regions without assigned peptides appear as a grey amino acid backbone. **D)** Canonical amino acid sequences for each matched gene symbol were evaluated for ibrutinib targets, as indicated by any or all lineage NPARC test with p < 0.05, and the proportion of sequences was compared to the proportion in the detection background for the full experiment. Proportional tests with Yates’ continuity correction were performed according to (*66*).

To validate our approach, we first examined whether TPP coupled with MS proteomics was able to identify BTK, the primary target of ibrutinib. BTK was only identified in the RCH-ACV lysates, which is consistent with the lineage background of this cell line. We identified two BTK proteoforms (BTK_1 and BTK_2) (Figure 1B), and both were stabilized in the presence of ibrutinib (Supplementary Figure 1B). Although BTK_1 was the only BTK proteoform that met the pAdj < 0.05 NPARC test significance threshold, BTK_2 was significantly shifted based on a p < 0.05 threshold. BTK_1 contained several peptides derived from the ATP binding site (Figure 1C, Supplementary Figure 1D) that were not detected in BTK_2 proteoform, which suggests that modifications, conformational changes, or splice variants reduced relative representation of these peptides in the BTK_2 proteoform. We did not observe cysteine 481 (C481), the specific residue that is covalently targeted by ibrutinib, in either proteoforms. We used BTK results to calibrate significance levels for further result interpretation, and we proceeded to consider pAdj < 0.05 as likely thermally impacted, and p < 0.05 as plausibly thermally impacted. In addition to BTK, previous studies(*18*) have revealed that ibrutinib binds a wide range of proteins (Supplementary Table 2), and our analysis confirmed several of these (Supplementary Figure 1C).

Ibrutinib’s C481 BTK binding site is flanked by G and L residues, and this sequence has been profiled in other off-target studies and is known to be homologous in off-targets(*33*). Although GCL enrichment was not significant in all detected proteins, we confirmed that ibrutinib targets display G*L motif enrichment, with notable over-representation of amino acids that can be modified by phosphorylation flanked by G and L residues (Figure 1A, Figure 1D). Taken together, these results indicate that TPP coupled with MS-based proteomics was able to detect over 2500 proteoforms representing close to 2000 proteins. Overall, the fact that we identified BTK as well as other known off-targets of ibrutinib serves as internal validation of our strategy.

### Using proteoform data to detect effects of drug binding on protein-protein interactions and complex formation

In addition to inhibiting activity of a target, drug binding may also lead to conformational changes or allosteric effects that change how the target interacts with its binding partners(*39*). To identify cases where thermal profiles indicated disruption of protein complexes, we performed an over-representation analysis (ORA) to identify proteins that form complexes in the CORUM database(*40*). Using FDR-adjusted hits (pAdj < 0.05), we identified 12 complexes (Figure 2A, Supplementary Table 3). Among these complexes, three hits (CORVET, HOPS, class C VPS complex) (Figure 2A-B, Supplementary Figures 2-3) share common components and have known interconnected biological functions as membrane tethering complexes(*41*) and as coordinators of signal transduction(*42*). Of particular relevance, this signaling includes NF-kB and AP1, which are induced during immunostimulation(*42*), and may represent a possible secondary route towards maintaining ibrutinib drug effects. Another identified complex, NUMAC (Supplementary Figure 4), integrates chromatin remodeling and histone methylation and plays a role in fine tuning gene expression during heart, lung, and immune cell development(*43*, *44*). The critical histone modifying component of NUMAC, CARM1, has been proposed as a cancer target and its knockdown enhances antigen-induced proliferation and cytotoxicity in tumor infiltrating T-cells(*43*), which is also observed in ibrutinib treated patients(*45*, *46*). Among other affected complexes, the p21(ras)GAP-FYN-LYN-YES complex, the CD20-LCK-LYN-FYN-p75/80 complex and the BRAF-MAP2K1-MAP2K2-YWHAE complex (Supplementary Figures 5-7), also have functional links to immunostimulatory signaling.

**Figure 2.**
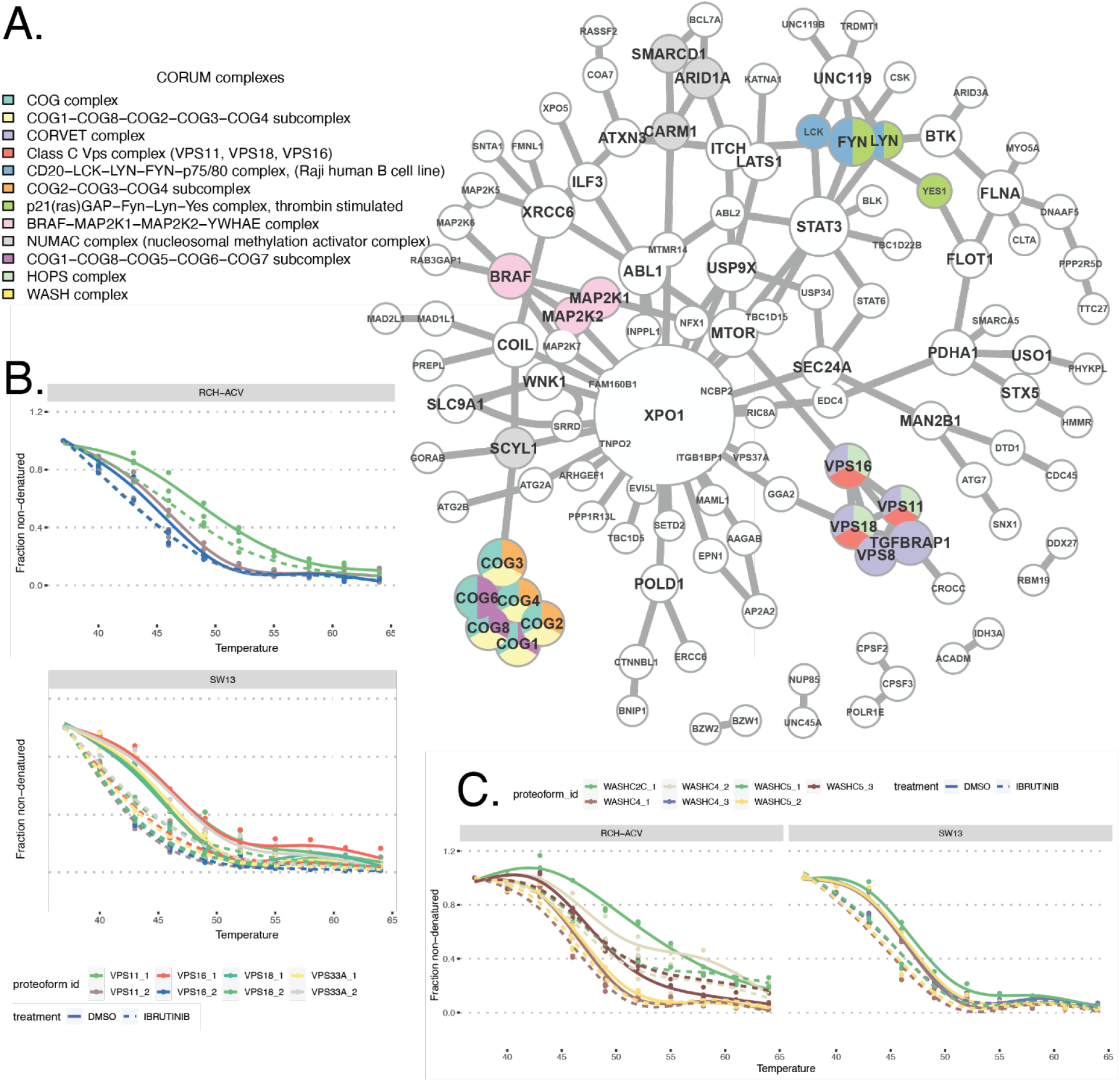
Physically associated and functionally implicated ibrutinib binding candidates: **A)** Network plot showing the CORUM complex composition of the sub-network of top NPARC hits and their associations according to the BioGRID interaction database. Nodes are plotted by size according to connectivity, and colored labels indicate membership in an enriched CORUM complex. **B)** Proteoform melting behavior for the HOPS complex, showing only results thresholded by individual cell line NPARC shifts with significance of p < 0.05. Note that the same proteoform colorings are plotted for each cell line, but that curves are filtered to show only significant proteoforms. **C)** Proteoform melting behavior for the WASH complex, showing only results thresholded by individual cell line NPARC shifts with significance of p < 0.05.

Protein-protein interactions are highly dependent on cell lineage(*47*), and we observed that the baseline thermal stabilities of complex-associated proteoforms and magnitude of drug-induced thermal changes were not uniform between cell lines. For example, the membrane tethering complexes were only indicated in SW13 (Supplementary Figures 2-3), and two of the immunostimulatory signaling complexes only in RCH-ACV (Supplementary Figures 5-6). To examine this further, we repeated the ORA tests within each cell line separately (Supplementary Table 3), which revealed several SW13-specific results including the multisynthetase complex, the EARP tethering complex, and the EIF2B2-EIF2B3-EIF2B4-EIF2B5 complex. These observations underscore that cell line-specific predominance of certain complexes could influence the range of interactions and scope of functional drug influence. Despite the cell type specificity of protein complex results, at proteoform-level, thermal changes still occurred for individual proteoform components across both cell lines, such as the HOPS complex (Figure 2B, Supplementary Table 1). This implicates individual proteoform targets from enriched complexes, even where the protein complexes themselves are not indicated.

Our analysis also identified complexes without clear functional links to the known on-target effects of ibrutinib. For example, the COG complex is essential in intra-Golgi transport and glycosylation of proteins and lipids(*48*)), and components were significantly thermally impacted by treatment in both cell lines (Figure 2A, Supplementary Figure 8). This complex is primarily found as an octamer in the cytosol, and components are also integrated into numerous other subcomplexes(*49*), which were also detected in the protein complex over-representation analysis (Supplementary Table 3). Although the link between ibrutinib and COG components is not known, changes in immunoglobulin glycosylation and secretion have been identified in ibrutinib treated CLL patients(*50*), in agreement with our results.

Another hit without a clear functional link to on-target pathways was the WASH complex (Supplementary Figure 9). This complex facilitates endosomal trafficking, surface receptor recycling(*51*) and maintenance of phagocytosis(*52*). Ibrutinib treatment is reported to cause defects in both receptor recycling(*53*), and endosomal trafficking(*54*, *55*), particularly of importance in mechanisms of aspergillosis susceptibility and immune cell egress. In addition to these potential links, the WASH complex modulates platelet function through reducing αIIbβ3 integrin cell surface expression(*56*), which mirrors a BTK-independent ibrutinib effect on αIIbβ3 integrin cell surface levels(*57*).

To further map the pathways affected by ibrutinib, as indicated by perturbations at the proteoform level, we used the BioGRID protein-protein interaction database(*58*) and performed functional enrichment analysis, using network topology analysis with network retrieval prioritization(*59*) (Supplementary Figure 10, Supplementary Table 4). Input hits were gene symbol IDs with at least one proteoform hit below the pAdj < 0.05 threshold for at least one cell type. As expected, we identified the B cell receptor signaling target pathway, in alignment with the intended function of ibrutinib; however, we also identified that ibrutinib had BTK-independent effects, including on protein autophosphorylation, RNA localization, and cell adhesion. Consistent with the results of our ORA analysis, other top hits included organelle membrane fusion and Golgi organization. Collectively, these analyses showcase how proteoform analysis of drug effects can reveal specific changes at the level of protein-protein interactions and complex formation. In the specific case studied here, observed effects cast new light on the potential, BTK-independent link between ibrutinib, and COG and WASH complexes, and their respective cellular functions.

### Proteoform analysis enables more nuanced target identification

Although we observed good agreement between our studies and previously reported ibrutinib off-target identification using kinobeads(*18*) (Supplementary Table 2), we also observed some additional hits, including several examples of clinically highly relevant targets. For example, although not seen in the kinobead study(*18*), we observed that BRAF had two proteoforms in our dataset (Figures 3A-B) and was also a component of a complex flagged in our ORA analysis (Supplementary Figure 7). One proteoform was a significant hit in both cell lines individually and across the entire dataset (Supplementary Table 1); however, the other, more thermally stable proteoform did not exhibit significant changes upon ibrutinib treatment for either cell line (Figure 3B). When BRAF results are taken in aggregate on a gene symbol level, the results indicate that BRAF is stabilized overall (Figure 3C), which illustrates that aggregation of this kind could mask the more nuanced behavior captured by the proteoform-specific analysis (Figures 3A-B). Therefore, the proteoform-level analysis suggests that BRAF may exist in distinct forms, one that binds ibrutinib and one that does not.

**Figure 3.**
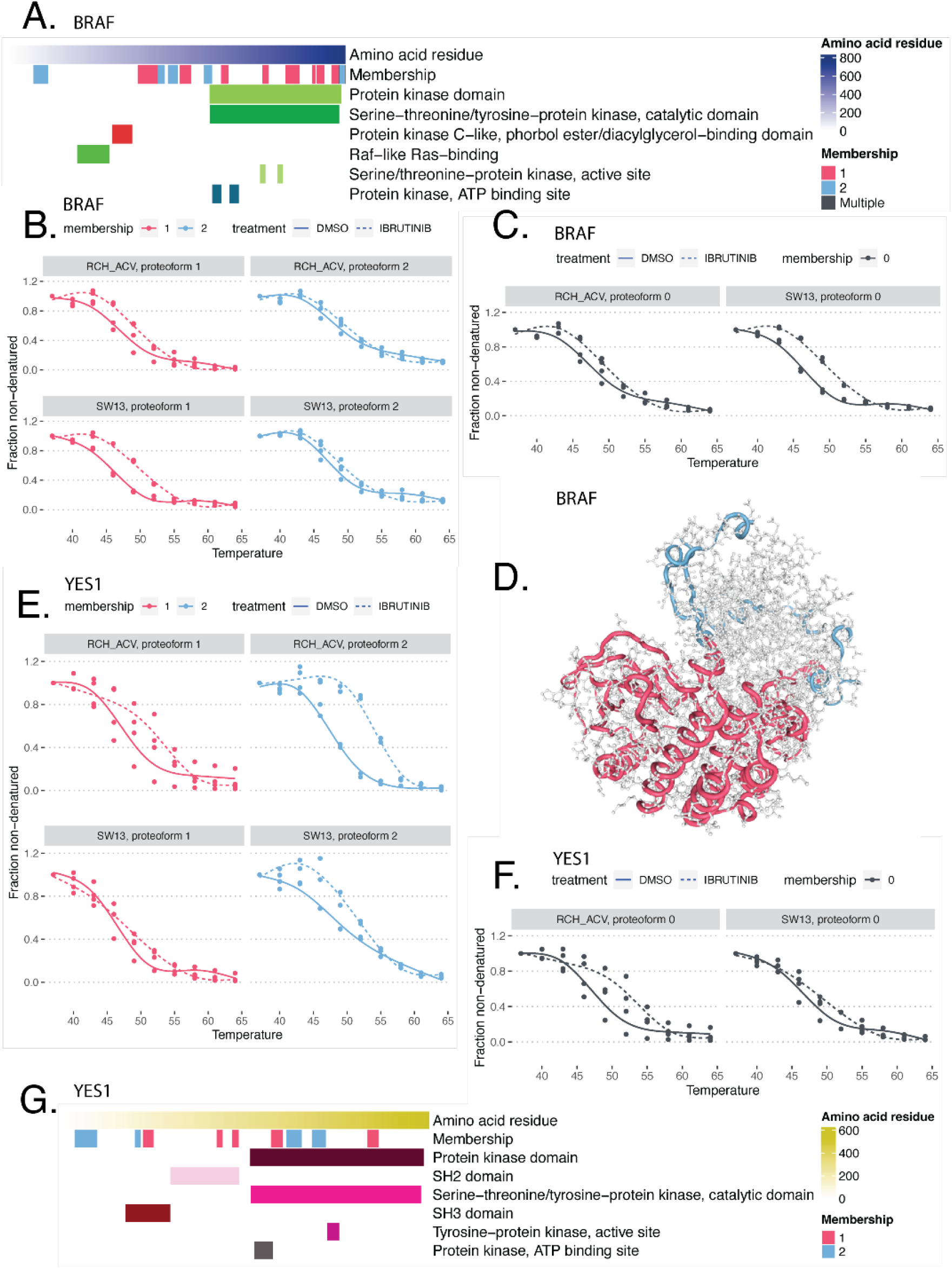
Specific ibrutinib target detection enabled by proteoform analysis: **A)** Peptides mapping to their corresponding locations on the canonical FASTA sequence for BRAF (sp|P15056|BRAF_HUMAN), colored by proteoform assignment “Membership” and highlighting regions with domains annotated in the interpro database **B)** Proteoform melting behavior for BRAF, separated and labeled by cell line, treatment, and proteoform. **C)** Protein aggregated melting behavior for BRAF, separated and labeled by cell line and treatment. **D)** Structural diagram of peptide mappings, plotted over the PDB structure for BRAF bound to MEK and an ATP analog, generated using the structure as published in (*61*), showing tube overlays for peptides colored by their proteoform assignments. Regions without assigned peptides or belonging to the MEK structure appear as a grey amino acid backbone. **E)** Proteoform melting behavior for YES1, separated and labeled by cell line, treatment, and proteoform. **F)** Gene symbol aggregated melting curve for the YES1 protein, separated and labeled by cell line and treatment. **G)** Interpro domains and associated peptide mappings for YES1 proteoforms.

Here, several lines of evidence support this possibility. On one hand, ibrutinib has been investigated for resensitizing BRAF-inhibitor refractory melanomas(*60*), a benefit not replicated with other BTK inhibitors indicative of ibrutinib-specific BRAF effects. Additionally, CORUM over-representation results indicated that ibrutinib treatment may interrupt BRAF interaction with MEK proteins (Figure 2A, Supplementary Figure 7), and experimental structure of the BRAF kinase domain in complex with MEK(*61*) appeared to be consistent with the mapping of BRAF proteoform 1 peptides (Figure 3D). On the other hand, the second proteoform may represent a dimerized form of BRAF. It is well established that BRAF is activated by RAS-dependent dimerization(*62*), including with other RAFs (ARAF and CRAF (RAF1)). Our results showed that RAF1 and ARAF were not thermally impacted in any sample or proteoform, leading us to propose that the proteoform of BRAF that is insensitive to ibrutinib are the hetero- or homodimers (Supplementary Figure 11 A-B). Furthermore, dimerized BRAF may be the main population of BRAF in cells, potentially explaining why kinobeads did not identify BRAF as ibrutinib target(*18*).

The identification of results with mixed drug binding affinity between proteoform subpools suggests that these gene symbol IDs would be harder to replicate in a traditional analysis. To probe this generalization further, we examined replication across the full dataset. Among the 217 pAdj < 0.05 gene IDs that were not replicated in the kinobeads study, 8.3% were hits for all proteoforms, with the rest having a proteoform above p = .05 or not clustered into proteoforms. But among the 12 kinobeads replicated gene IDs, 42% were clustered into proteoforms and were significant for all proteoforms. This demonstrates that results previously identified at gene symbol level(*18*) were proportionately ∼5x more likely to be thermally impacted for all clustered proteoforms, compared to unreplicated gene symbols.

Thermal proteome profiling may generally be a more sensitive approach for certain drug contexts. However, not all replicated hits from the kinobeads study would have been clear without proteoform clustering. For example, we identified two proteoforms of a tyrosine kinase YES1, an important member of the src kinase family. YES1_2 was stabilized in RCH-ACV and across the whole dataset, but YES1_1 was not (Figure 3E). Without proteoform clustering, in the gene symbol aggregated full dataset, YES1 fell outside the p-value threshold at p = 0.0617, pAdj = .433 (Figure 3F). Structurally, the stabilized YES1_2 peptides mapped in the N-terminal region, which in src kinases is known as the src N-terminal regulatory element (SNRE), an understudied and intrinsically disordered region thought to perform lipid binding, enacting regulatory functions and enabling cell type-specific roles(*63*) (Supplementary Figure 11D). These context-dependent lipid interactions could be another mechanism introducing melting variance between samples. The apparent baseline melting difference between RCH-ACV and SW13 is notable, despite a lack of mutations in this protein detected in the depmap mutation profiling dataset(*36*), further supporting that YES1 baseline variation occurs and can be independent of genetic sequence. Together, this indicates the YES1 protein could be susceptible to a range of important conformational states or interaction partners that affect proteoform-level thermal stability between cell lines, and which could also limit or enable context-dependent ibrutinib binding.

Collectively, these examples illustrate how proteoform-level analysis could enable a more nuanced interpretation of drug binding, one that is highly context and cell line dependent. Additionally, comparing and contrasting results obtained at the aggregate gene symbol level and proteoform level may indicate functionally relevant differences worth examining further.

### Validation by peptide resolved pulldown experiments

To further validate the ability of our proteoform-level drug binding approach to identify relevant drug-proteoform interactions, we performed an ibrutinib pulldown using the RCH-ACV cell line. Results were considered significant if they were replicated in two out of three ibrutinib treated preparations without DMSO detection, and results detected in both ibrutinib and DMSO pulldown samples were considered if they were replicated in at least two preparations and also had intensities that met a paired t-test threshold of p < 0.05. The pulldown method has several notable limitations, namely that it detects only more stable, direct drug interactions rather than weaker interactions or those associated with protein complex effects, and this method is additionally limited by its high peptide detection threshold. Despite these considerations, we could nevertheless confirm 7 proteoform-resolved pulldown hits from the 27 detected from peptide-matched RCH-ACV TPP proteoform hits, including multiple novel results (Figure 4A; Supplementary Table 5). Among proteoform results that were replicated in the pulldown, we identified the WASH complex component WASHC2C_1 (Figure 4A-B), and both proteoforms of BTK (Figure 4C). We observed that WASHC2C_1 had higher mean pulldown intensity than the BTK proteoform BTK_2 (WASHC2C_1 = 5.64 x 10^6^, BTK_2 = 7.48 x 10^5^), although it was only detected at the 20uM ibrutinib probe dose. The second proteoform that was not thermally impacted, WASHC2C_2, was also not detected in the pulldown (Figure 4B). This preference of ibrutinib for WASHC2C_1 could hint at differential post-translational modifications, lipid interactions, or other structural differences between these proteoforms that might influence binding affinities. Alternatively, these differences might indicate that WASHC2C_1 is a more prevalent form in the RCH-ACV cell line, which also has implications for ibrutinib targeting. Taken together, our study indicated that TPP MS-based proteomics coupled with proteoform detection can lead to comprehensive identification of unknown targets; more specifically, our results indicate that the relationship between WASH complex and ibrutinib is likely functionally relevant and requires further study.

**Figure 4.**
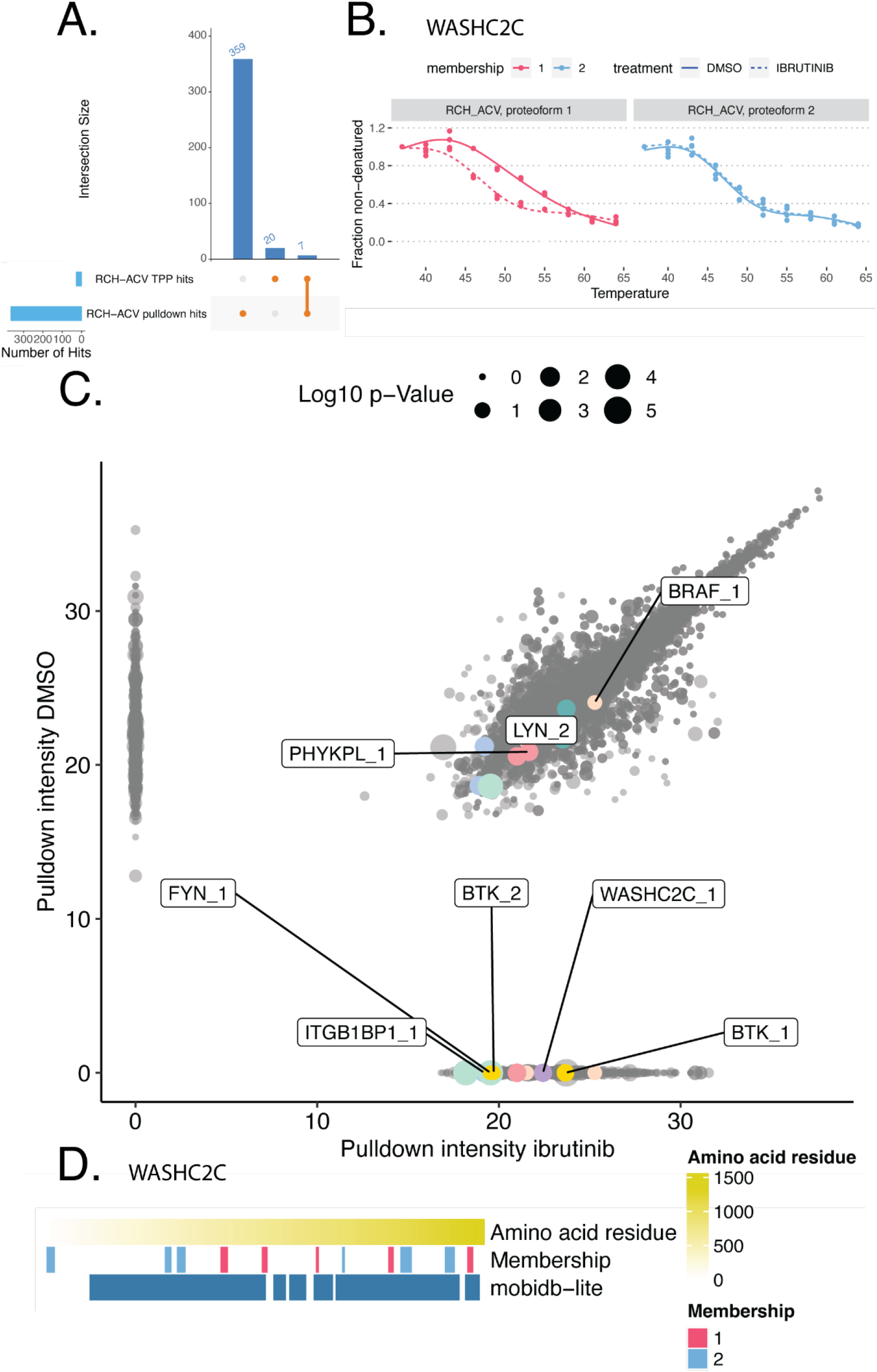
Ibrutinib target validation by pulldown with proteoform aggregation: **A)** Detected hits (p < 0.05) in each individual cell line stratified by their gene symbol ID, and showing their overlap with the previously identified functional ibrutinib target hits. **B)** Proteoform melting behavior for WASHC2C in RCH-ACV, separated and labeled by treatment and proteoform. **C)** Pulldown intensities for each replicate and dose, adjusted by log2(intensity + 1). Point sizes indicate pAdj in the RCH-ACV TPP dataset, as obtained using NPARC to distinguish treated and DMSO melting. **D)** Mobidblite mappings for WASHC2C peptides by proteoform, representing disordered regions.

## Discussion

This work demonstrates the potential of proteoform-level deconvolution to identify new targets of drugs, using ibrutinib as a case study. Proteoforms can be identified from untargeted thermal proteomics data and in relevant cellular contexts by applying our previous methods(*23*), and here we illustrate that proteoforms can also be distinguished with respect to their drug binding abilities, leveraging the thermal impact of a drug for treatment-specific proteoform identification. This enables deeper interpretation of the functional implications of drug activity, and paves the way for identification of specific proteoforms and the roles they perform. In turn, we expect that this will further expand the range of therapeutically relevant targets and improve the precision of personalized medicine. Moreover, these types of studies will improve our understanding of adverse side effects, mechanisms of action, mechanisms of resistance, and polypharmacology. For example, our study highlights the possibility that ibrutinib impacts many more off-targets than previously known, which may converge in their functional roles and impacted pathways. These results have far-reaching implications in interpreting the primary and secondary effects of ibrutinib treatment. For instance, newly identified ibrutinib-impacted proteoforms play roles in mechanisms that may amplify drug efficacy, such as B-cell receptor signaling, induction of bone marrow egress, and T-cell immunomodulation. We also uncovered other functions potentially directly impacted by ibrutinib, such as Golgi trafficking, glycosylation, and cell adhesion. Collectively, the extended target list could enable elucidation of the complexities of BTK-independent ibrutinib immunomodulation. Additionally, these results collectively provide context for understanding and addressing multi-causal common effects of clinical importance, such as defects in anti-fungal immunity(*55*) Aspergillosis is a very clinically common secondary infection (*64*) with unclear etiology to explain its high incidence, but which has been previously linked to endosomal defects (*54*, *55*) in addition to immunosuppression. Together, these results provide a new foundation for examining and responding to repurposing indications and treatment toxicities.

Although this approach extends the capabilities of proteoform-specific drug target deconvolution, it has several limitations that are worth noting. In general, our proteoform identification requires in-depth peptide detection, as well as significant instrument time and resources, which could potentially be improved by method optimization. More specifically, although our experiments captured a number of previously reported ibrutinib targets, we were not able to confirm all of them, most notably TEC kinase family members(*65*). Therefore, tailoring additional experimental setups to capture more realistic cellular or tissue environments may be needed to address additional targets of drug activity.

Moving forward, we believe that our results and methods offer valuable insights for the refinement of preclinical research strategies and rationalize clinical observations. Continued research that extends these methods and incorporates additional therapeutic agents, cellular contexts, and functional interpretation approaches could enhance our understanding of drug mechanisms and lead to better tailored precision medicine approaches.

## Supplementary Figures

**Supplementary Figure 1.**
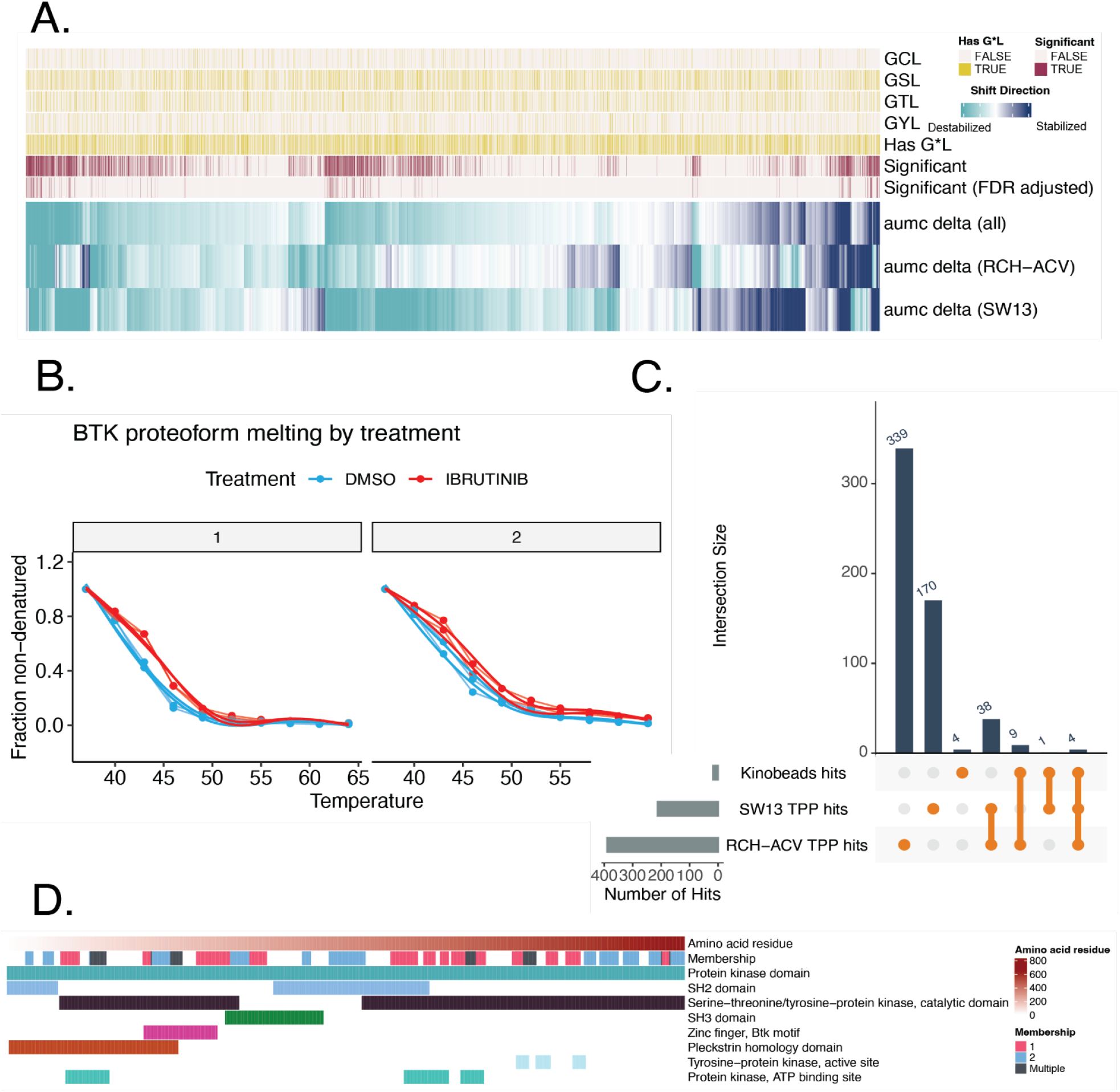
Contextual details of ibrutinib binding targets and BTK proteoforms: **A)** All proteoform results for which full melting curve models could be generated are plotted clustered by delta aumc, defined as the aumc of DMSO data with aumc of ibrutinib data subtracted. Highlighted proteoforms are annotated by presence of GCL, GSL, GTL, or GYL sequences, and by their significance in one or both cell line(s) at a threshold of p < 0.05 “Significant” or pAdj < 0.05 “Significant (FDR adjusted)”. **B)** Fraction non-denatured for each BTK proteoform, demonstrating melting of the four replicate curves by treatment. **C)** Detected hits (p < 0.05) in each individual cell line stratified by their gene symbol ID, and showing their overlap with the previously identified functional ibrutinib target hits from (*18*). **D)** Peptides mapping to their corresponding locations on the canonical FASTA sequence for BTK, colored by proteoform assignment “Membership” and highlighting regions with domains annotated in the interpro database (*67*).

**Supplementary Figure 2.**
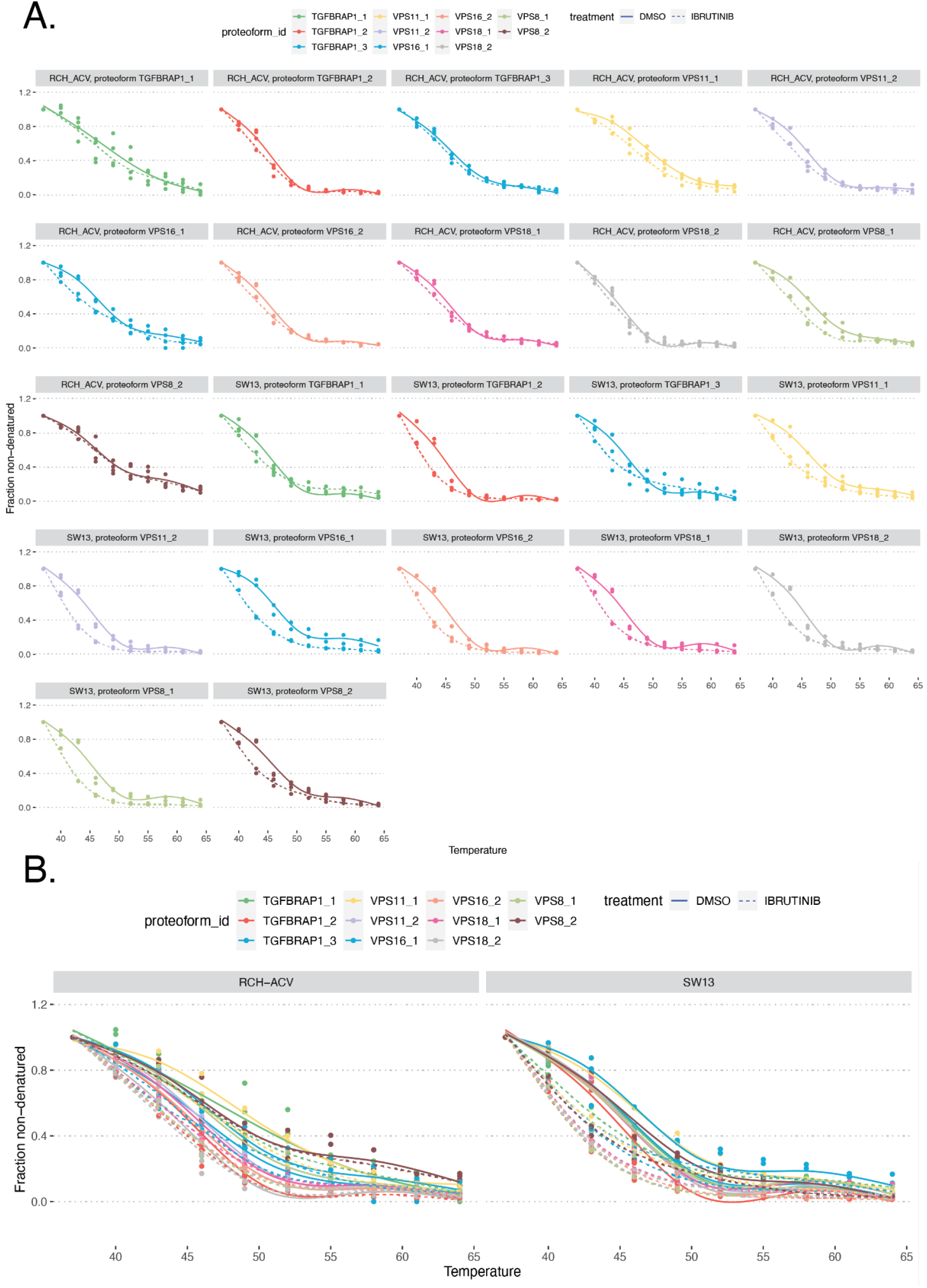
Proteoform composition and thermal behavior of the CORVET complex: **A)** All proteoform melting, for the CORVET complex. **B)** Same data as (A), plotted in one window per cell line

**Supplementary Figure 3.**
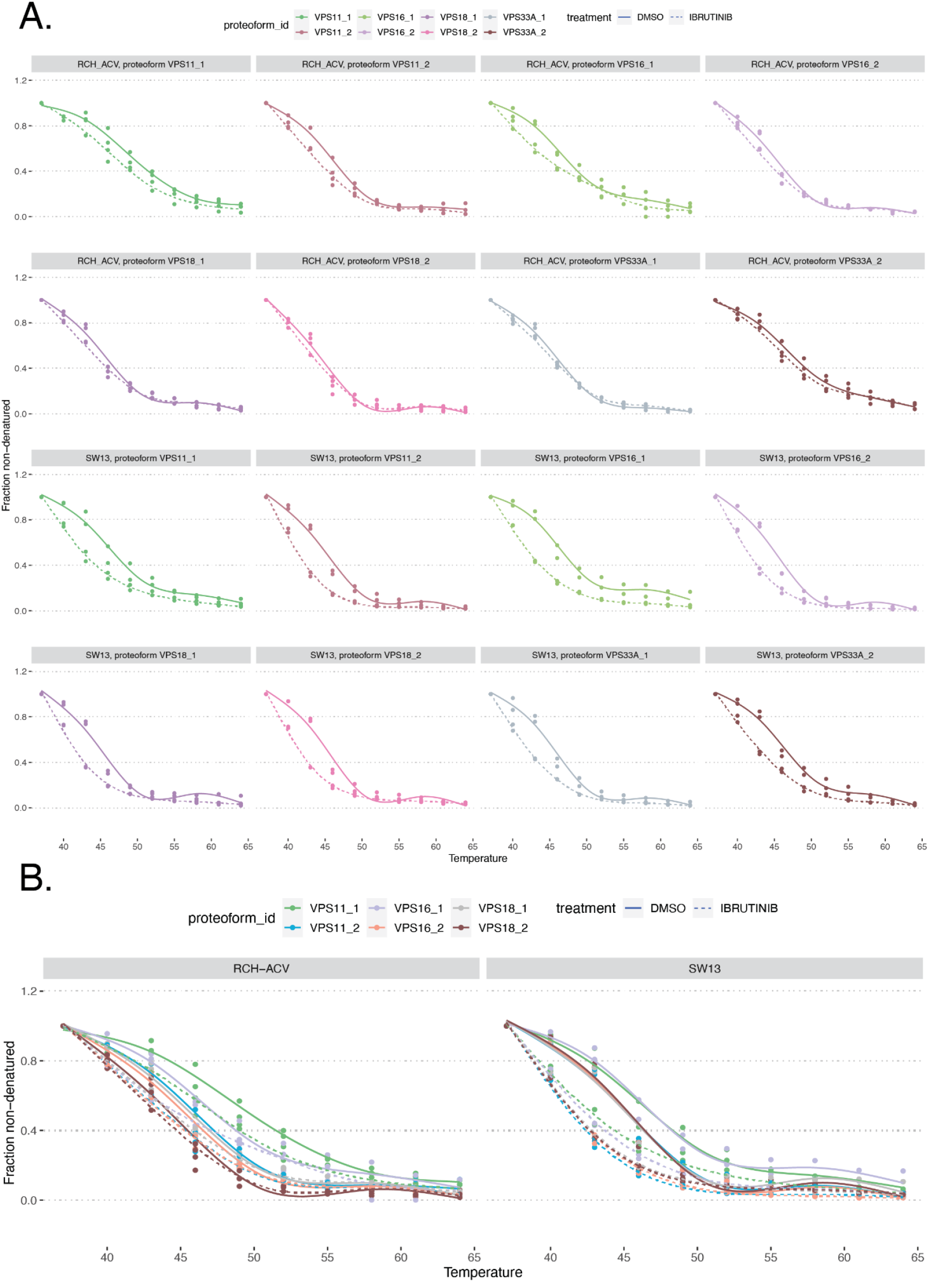
Proteoform composition and thermal behavior of the HOPS and class C VPS complex: **A)** All proteoform melting, for the shared components of the HOPS and class C VPS complexes. **B)** Same data as (A), plotted in one window per cell line

**Supplementary Figure 4.**
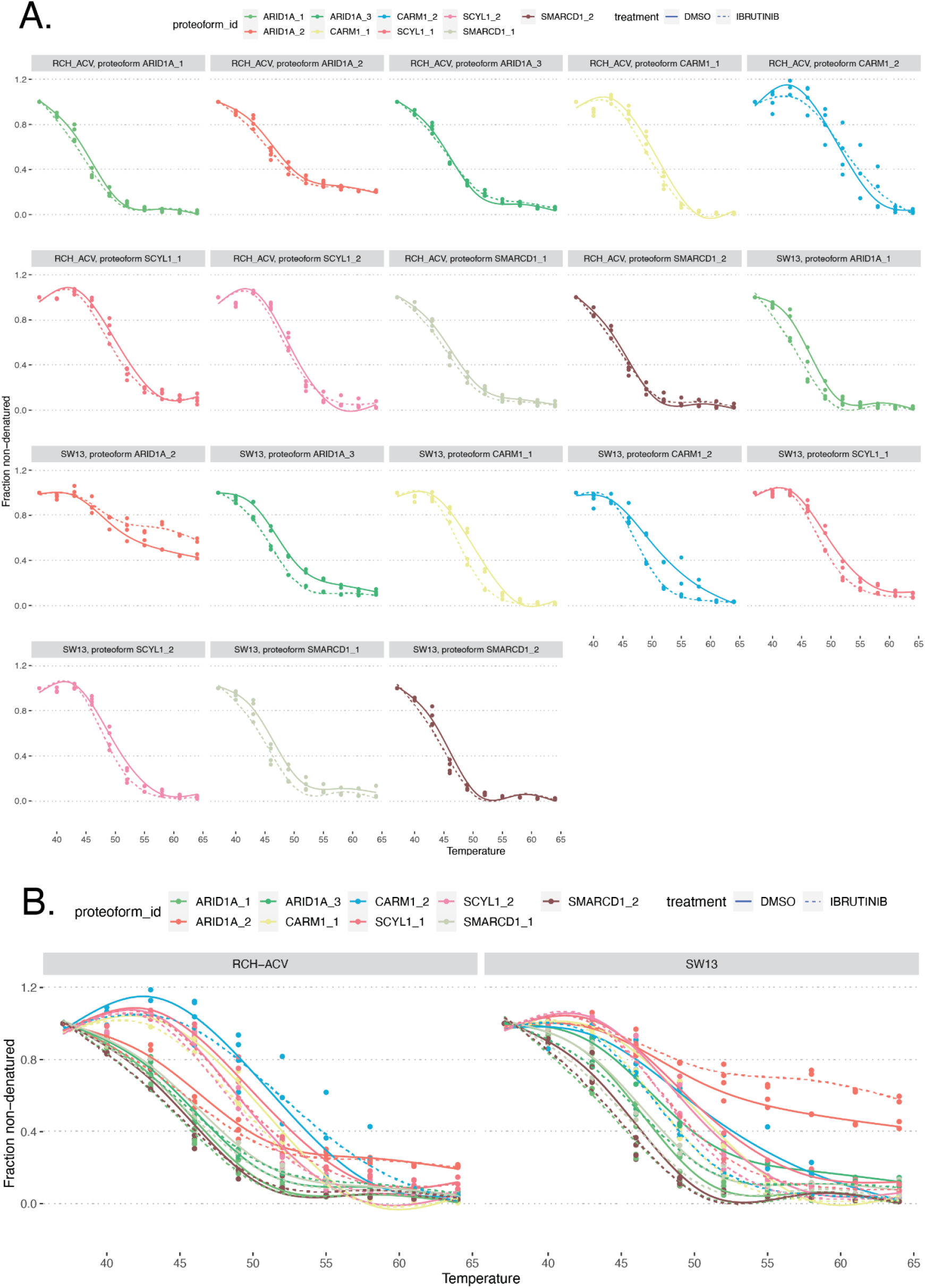
Proteoform composition and thermal behavior of the NUMAC complex: **A)** All proteoform melting, for the NUMAC complex. **B)** Same data as (A), plotted in one window per cell line

**Supplementary Figure 5.**
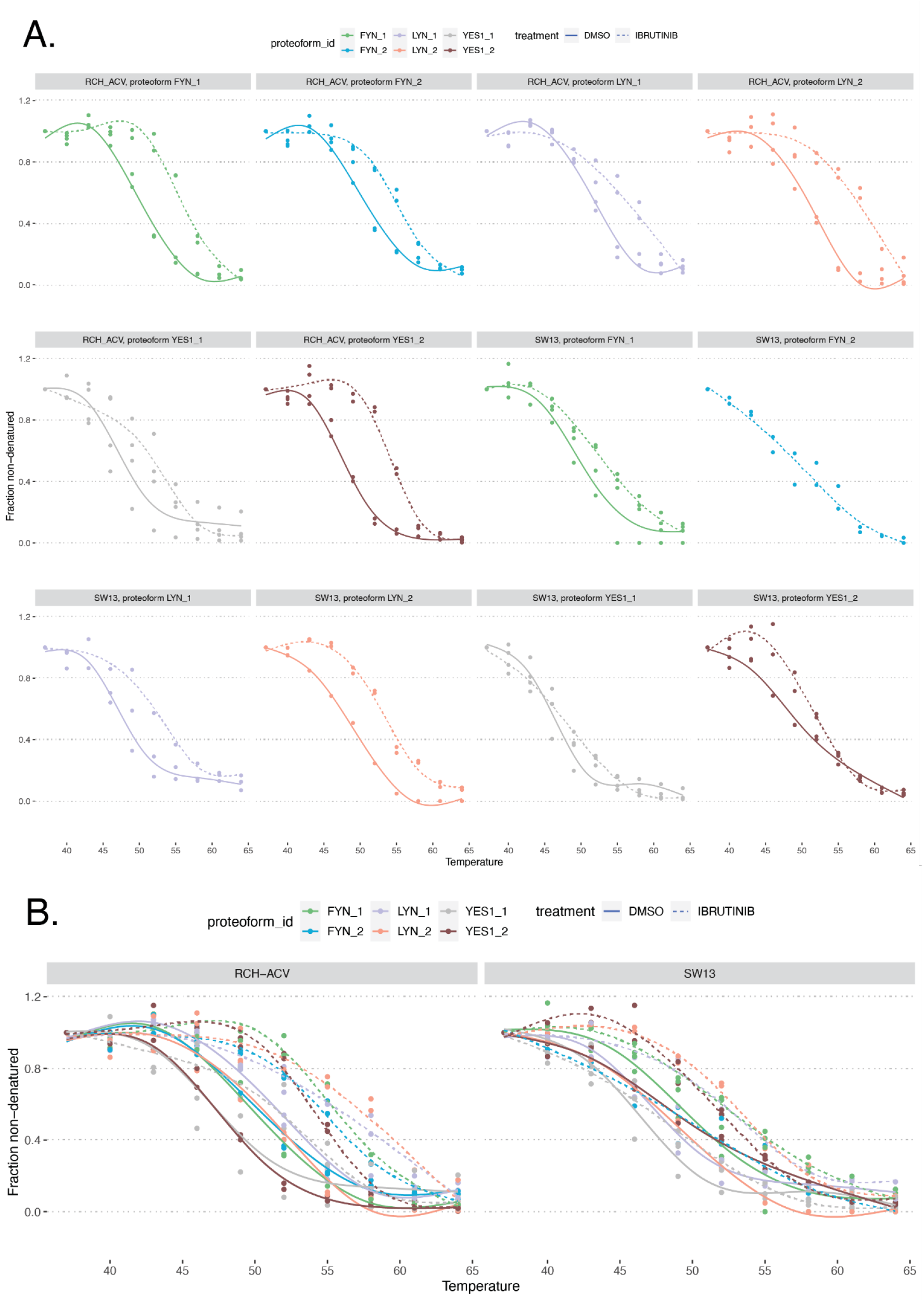
Proteoform composition and thermal behavior of the p21(ras)GAP-FYN-LYN-YES complex: **A)** All proteoform melting, for the p21(ras)GAP-FYN-LYN-YES complex. **B)** Same data as (A), plotted in one window per cell line

**Supplementary Figure 6.**
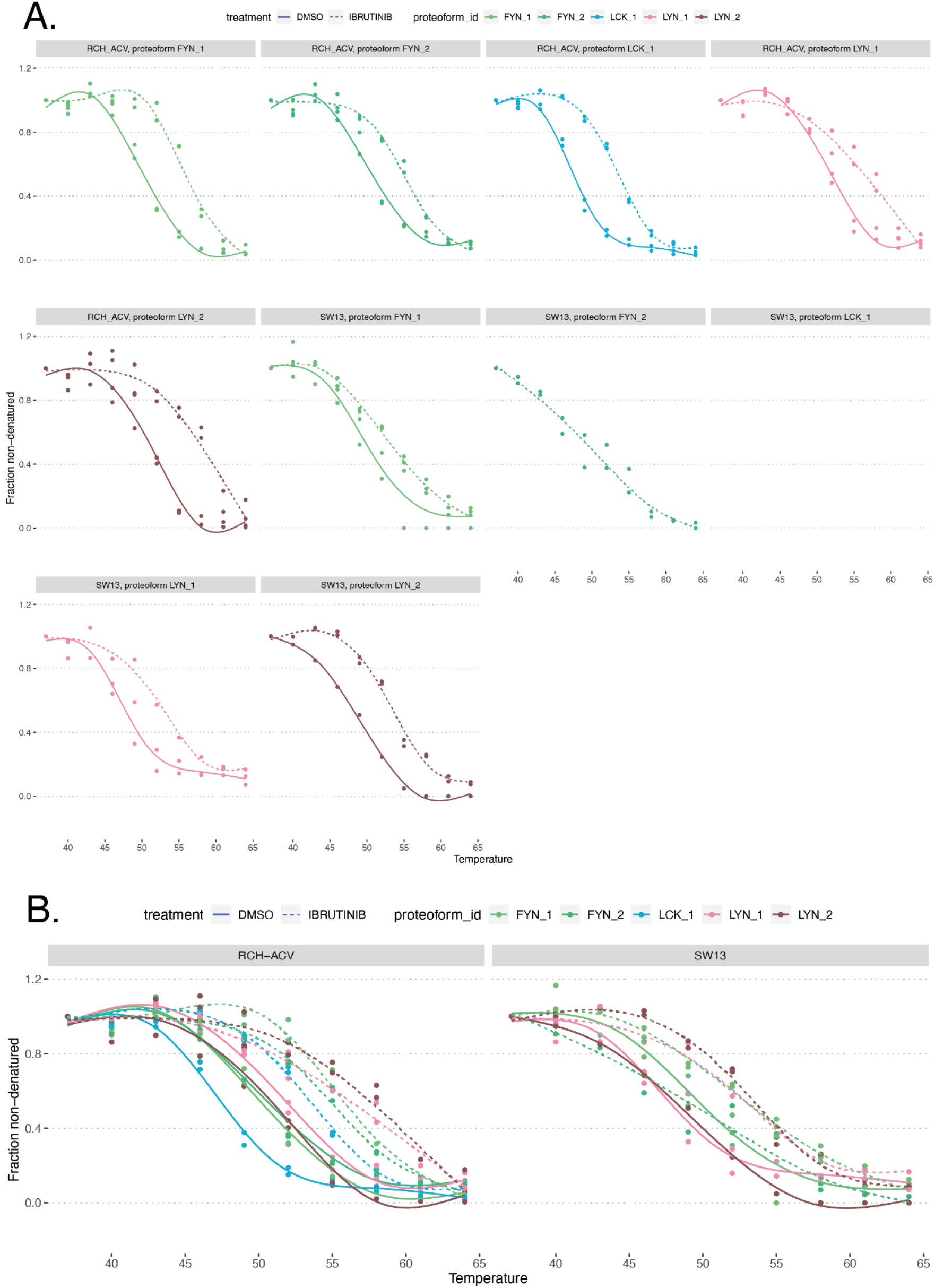
Proteoform composition and thermal behavior of the CD20-LCK-LYN-FYN-p75/80 complex: **A)** All proteoform melting, for the CD20-LCK-LYN-FYN-p75/80 complex. **B)** Same data as (A), plotted in one window per cell line

**Supplementary Figure 7.**
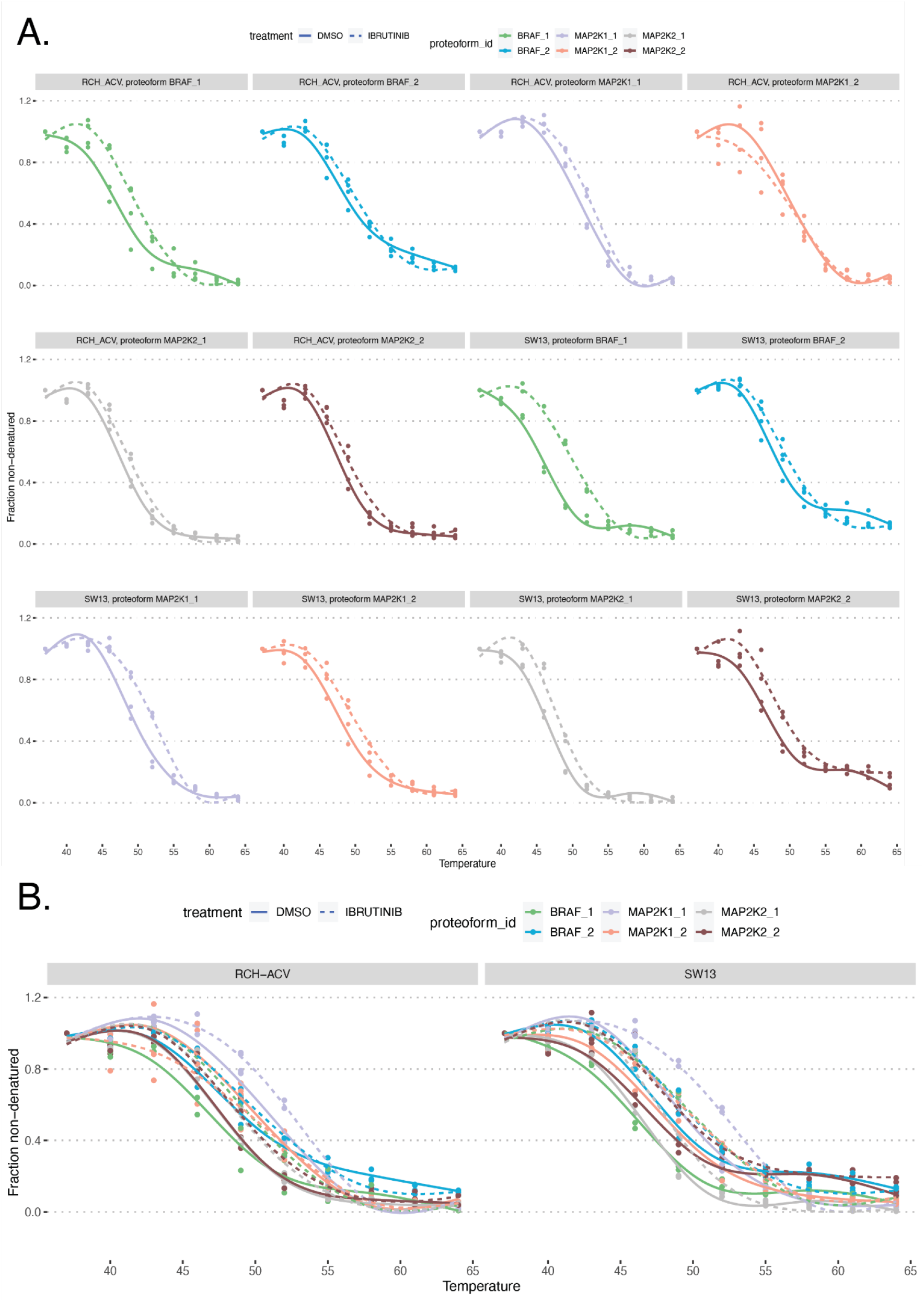
Proteoform composition and thermal behavior of the BRAF-MAP2K1-MAP2K2-YWHAE complex: **A)** All proteoform melting, for the BRAF-MAP2K1-MAP2K2-YWHAE complex. **B)** Same data as (A), plotted in one window per cell line

**Supplementary Figure 8.**
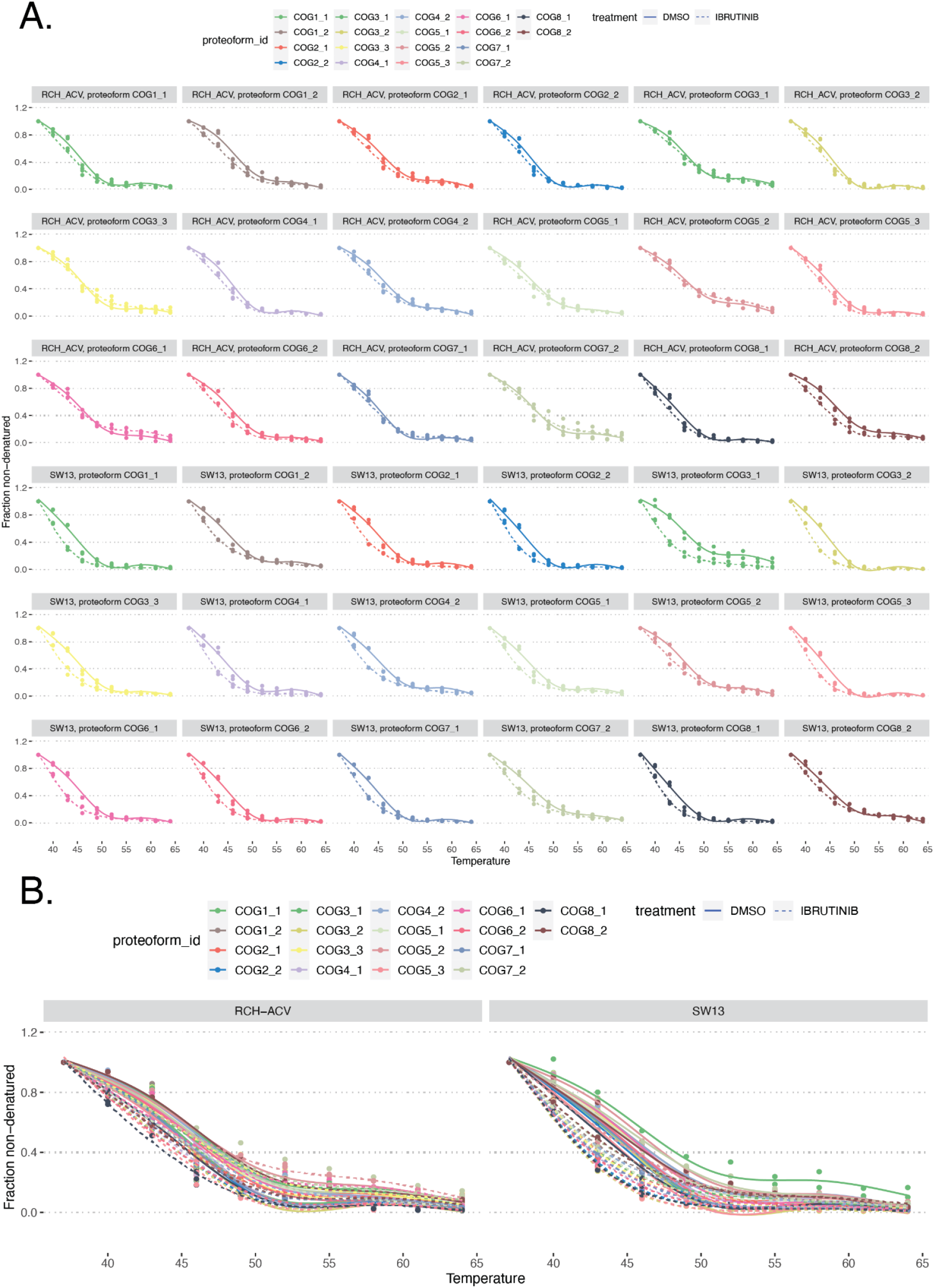
Proteoform composition and thermal behavior of the COG complex: **A)** All proteoform melting, for the COG complex. **B)** Same data as (A), plotted in one window per cell line

**Supplementary Figure 9.**
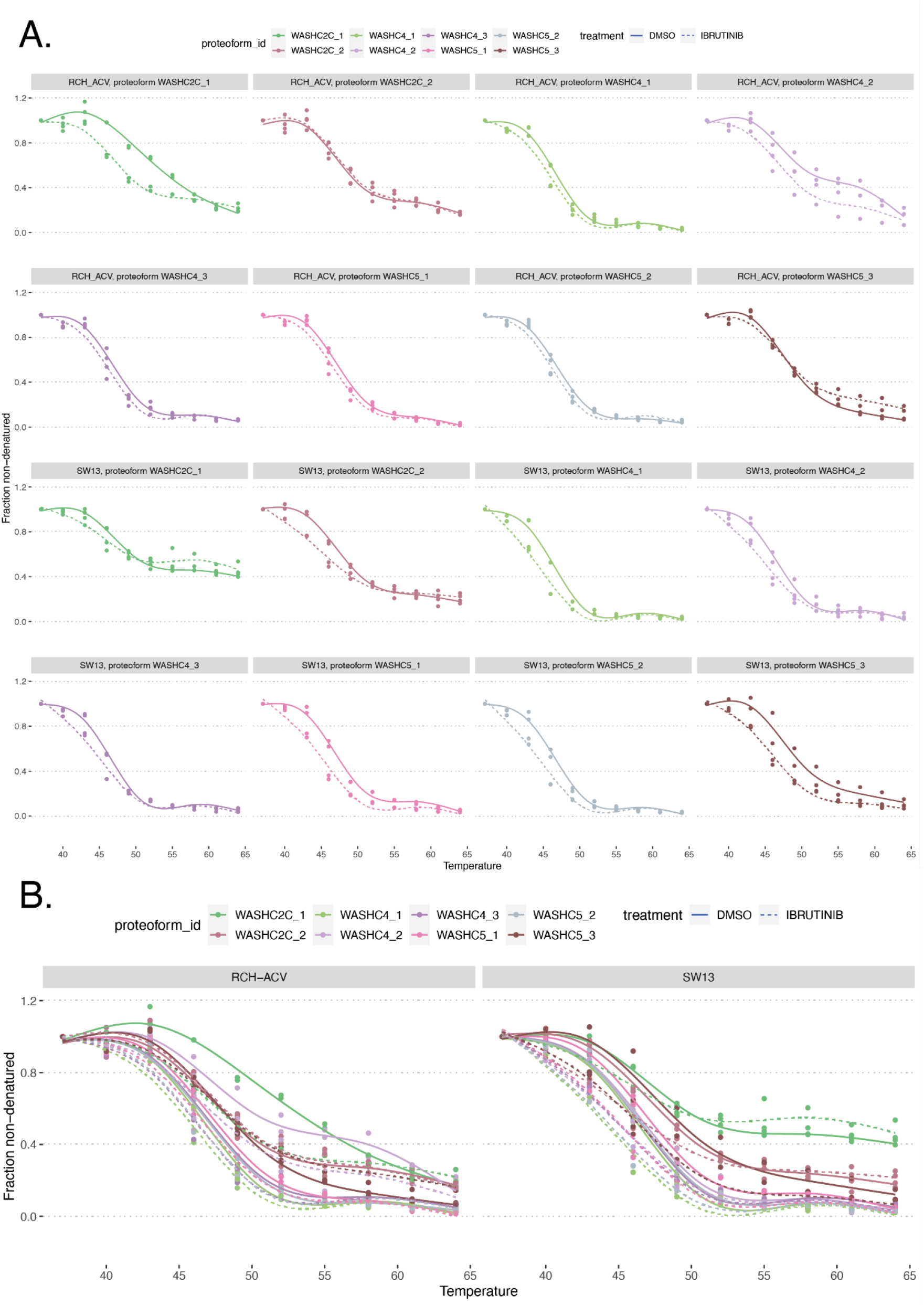
Proteoform composition and thermal behavior of the WASH complex: **A)** All proteoform melting, for the WASH complex. **B)** Same data as (A), plotted in one window per cell line

**Supplementary Figure 10.**
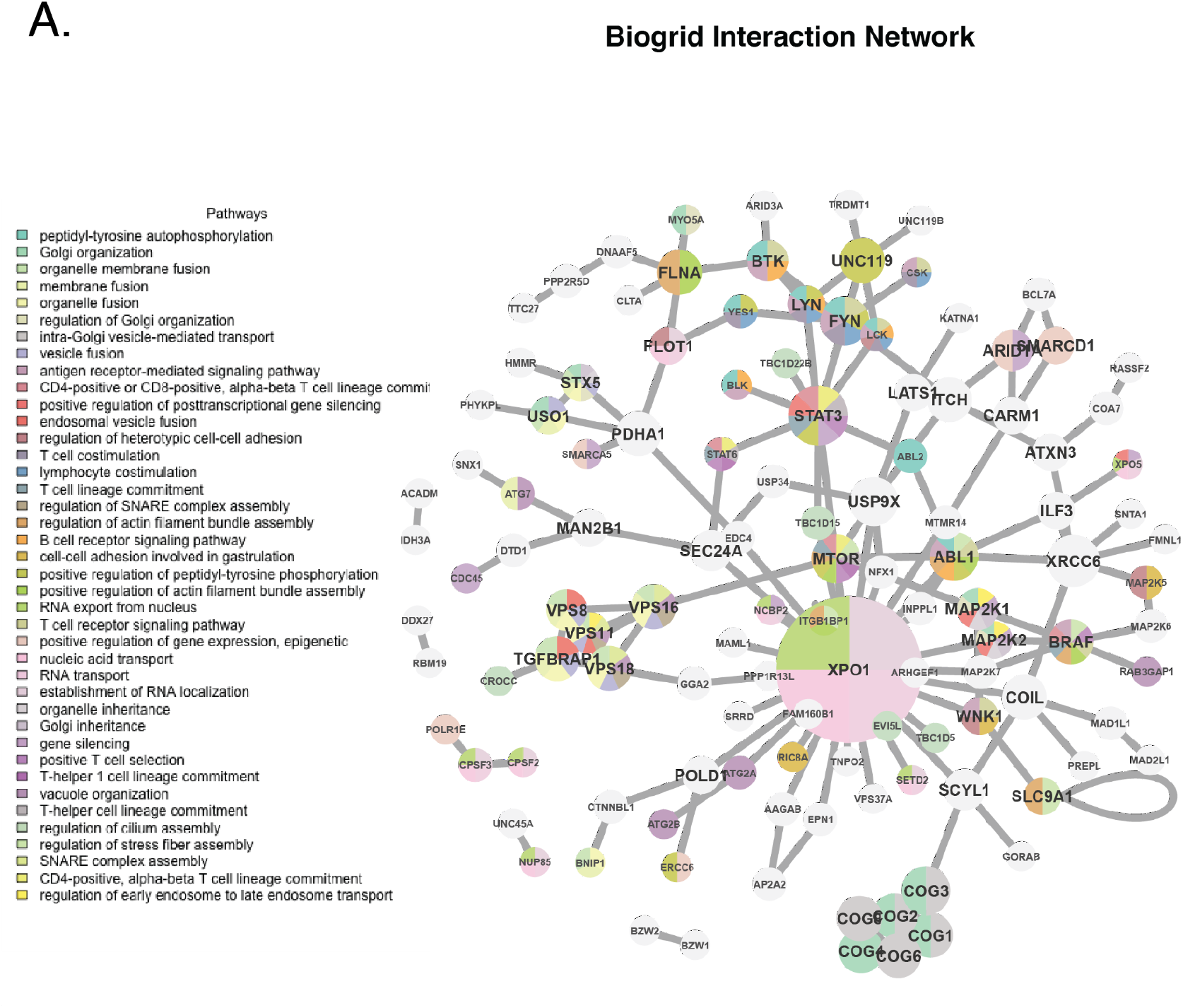
Pathway representation in interaction network analysis: **A)** Network plot showing the GO:biological process pathway distribution, plotted over the sub-network of top NPARC hits and their associations according to the BioGRID interaction database. Nodes are plotted by size according to connectivity, and colored labels indicate membership in an enriched GO:biological process pathway.

**Supplementary Figure 11.**
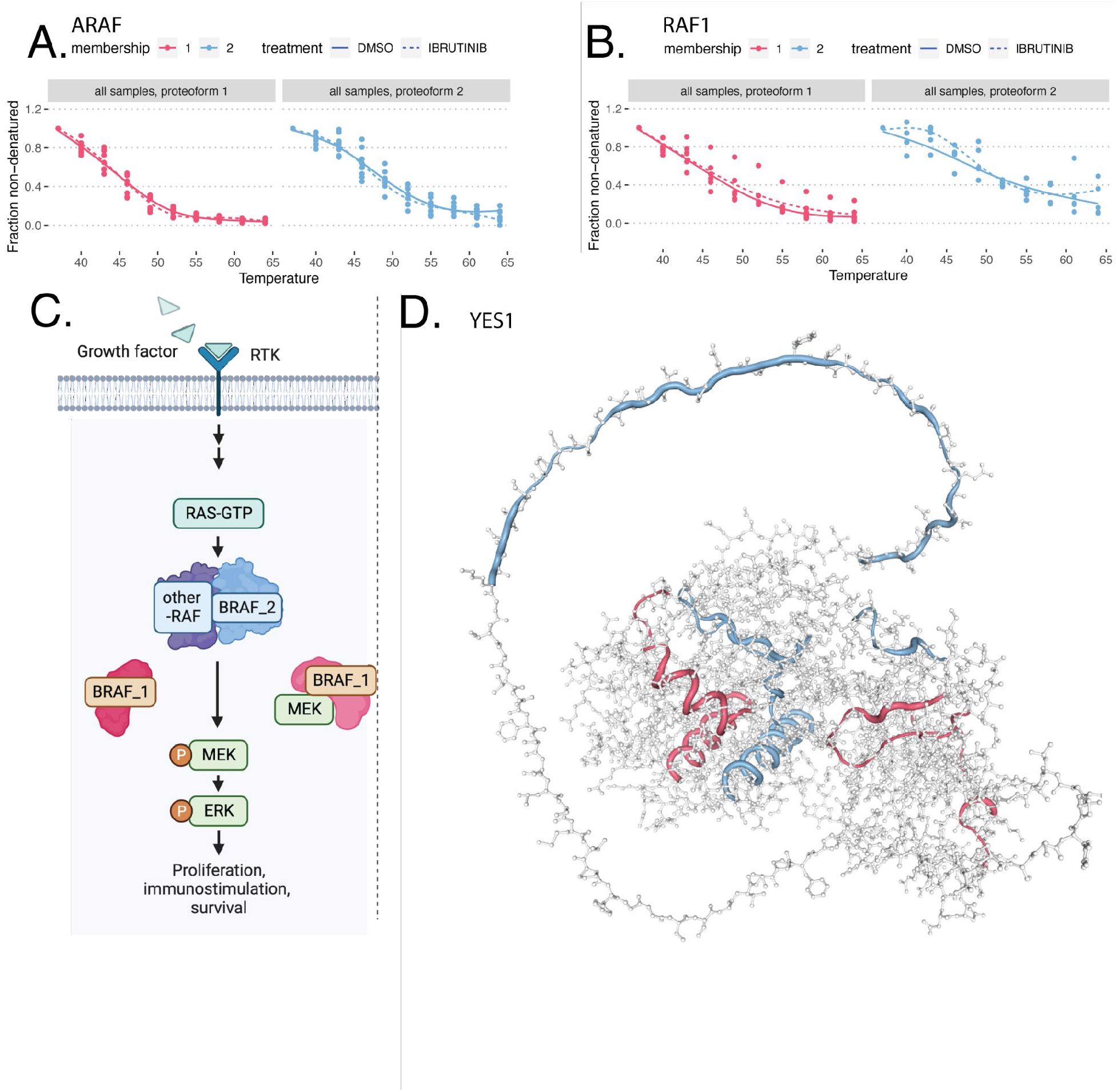
Ibrutinib target refinement: **A)** Proteoform melting behavior for RAF dimerization partner ARAF, showing data from all cell lines separated by proteoform. **B)** Proteoform melting behavior for the single proteoform detected for RAF dimerization partner RAF1, showing data from all cell lines. **C)** Illustrated diagram of proposed BRAF proteoforms, created with BioRender. **D)** Alphafold structure for YES1 generated using the canonical FASTA sequence (sp|P07947|YES1_HUMAN), showing colored tube overlays for peptides colored by their proteoform assignments. Regions with multiple matched proteoforms are displayed with blended translucent coloring, and regions without assigned peptides appear as a grey amino acid backbone.

## Materials and methods

### Cell lysis and protein concentration

RCH-ACV or SW13 cell pellets were thawed on ice, and resuspended in 100 µL of a HEPES buffer (10 mM HEPES, 20 mM MgCl2). Cells were subjected to three freeze-thaw cycles in liquid nitrogen and a 37°C water bath, respectively, followed by mechanical disruption via syringe inversion. Protein concentration was determined using the DC protein assay (Bio-Rad) according to manufacturer specified instructions.

### Desalting of peptides

Desalting was performed using 100 mg SepPak cartridges. Prior to use, cartridges were conditioned by first wetting them with 1 mL of 100% methanol (MeCN), followed by a second wetting step using 1 mL of an 80% MeCN solution containing 0.5% formic acid (FA). Next, cartridges were equilibrated by passing through 3 mL of 0.1% trifluoroacetic acid (TFA). Subsequently, the samples were acidified to a pH range of 2-3 using formic acid before being loaded onto the cartridges. After sample loading, cartridges were washed and desalted by passing 3 mL of 0.1% TFA through them. This was followed by an additional wash with 0.25 µL of 0.5% FA. Peptides were then eluted from the cartridges using two 500 µL aliquots of 80% MeCN and 0.5% FA. Eluted peptides were subsequently dried using a speed-vac.

### Mass spectrometry proteomics

A 3-hour gradient on a Q-Exactive mass spectrometer was run with the following gradient profile: 0-6 min: 3% 12 min: 6% 185 min: 37% 190 min: 42% 192 min: 99% 200 min: 99% 203 min: 3% 213 min: 3%

Detection Settings: First scan: 70K resolution, 1e6 AGC target, 100 ms max IT, mass range 300-1600 m/z. Data-dependent MS/MS: 35K resolution, 1e5 AGC target, 150 ms max IT, Top5 selection, 2 m/z isolation window, 1e3 min AGC target, 30s dynamic exclusion.

Buffers Used: NanoA: 90% MeCN, 5% DMSO, 5% H2O, 0.1% FA NanoB: 95% H2O, 5% DMSO, 0.1% FA LoadA: 97% H2O, 3% MeCN, 0.1% FA LoadB: 95% MeCN, 5% H2O, 0.1% FA

### Mass spectrometry data search

Raw mass spec outputs were processed for quality control and quantified with a standardized pipeline, ddamsproteomics version 1.0.2, openly accessible at: https://github.com/lehtiolab/ddamsproteomics/releases/tag/v1.0.2.

The following adjustable parameters were specified: --genes --hirief --fractions --symbols --isobaric tmt10plex --denoms ‘DMSO1:126 DMSO2:126 IBRUTINIB2:126 IBRUTINIB1:126’

The mapping database was Homo_sapiens.GRCh38.92

### Lysate preparation and thermal proteome profiling

RCH-ACV or SW13 cell extracts were treated with either Ibrutinib or vehicle (DMSO) in duplicates for 10 min at 20°C and with gentle shaking at 700 rpm. The treated lysates were aliquoted, and the aliquots were heated at 10 designated temperatures ranging from 37 to 67°C in order to denature and aggregate the proteins. The remaining soluble protein fraction was cleared from the heat-aggregated proteins by means of centrifugation (40 minutes, 21000g, 4°C) and the resulting soluble protein fractions were prepared for LC-MS/MS analysis. Samples proceeded to digestion, desalting and mass-spectrometry proteomics data acquisition (see above). Additionally, samples were labeled using 10-plex TMT tags (Thermo Fisher Scientific), with labeled sets established using one 10-plex set for each corresponding 10 point melting curve and using the same amount of respective label for each sample. Labeling was performed according to the manufacturer’s instructions but with 2-hour incubation before quenching the TMT labeling reaction. Labeling efficiency was determined by LC–MS/MS before mixing the TMT-labeled samples. Prior to LC-MS/MS analysis, the tandem mass tagged peptides were pre-fractionated using high-resolution isoelectric focusing (HiRIEF) (*37*) as well as additional fractionation using reversed-phase chromatography at pH 12.

### Pulldown in lysate

To ensure the inference of binding results is appropriate, an ibrutinib pulldown was performed using the RCH-ACV cell line, with results performed in triplicate and in two probe concentrations. The lysate was treated with DMSO, 10 µM, and 2 mM of an Ibrutinib-probe. For each treatment, samples were incubated for 10 minutes at 20°C and with gentle shaking at 700 rpm. Afterward, lysates were centrifuged at 21,000g for 1 hour at 4°C. A premix containing TFL (10 mM in DMSO), TCEP (52 mM, 15 mg/mL in ddH2O), 1xTHPTA (1.667 mM in H2O), and CuSO4 (50 mM) was added to each sample. The samples were incubated at room temperature for 1 hour. Post-incubation, the samples were treated with cold acetone for protein precipitation, stored overnight at -20°C, and subsequently subjected to centrifugation at 21,000g for 15 minutes at 4°C. Proteins were pelleted and washed twice with cold methanol. The pellet was resuspended using a probe sonicator and treated with 0.2% SDS in DPBS with the addition of 0.6 M urea. Protein concentration was determined, and equal amounts of protein were transferred to Protein LoBind tubes containing prewashed (1 mL 0.2 % SDS in DPBS) neutravidin beads. Samples were incubated for 1 hour under continuous mixing, followed by washing with 0.2 % SDS in DPBS, 6 M urea in ddH2O, and DPBS. Following this step, samples proceeded to on-bead digestion, desalting and mass-spectrometry proteomics data acquisition. Prefractionation for this experiment was performed using high pH fractionation. The data search was performed as described above. Results were considered significant if they were replicated in two out of three ibrutinib treated preparations without DMSO detection, and results detected in both ibrutinib and DMSO pulldown samples were considered if they were replicated in at least two preparations and also had intensities that met a paired t-test threshold of p < 0.05.

### On-bead digestion

The beads were resuspended in a buffer containing 2 M urea in 50 mM TEAB and subjected to reduction and alkylation steps with 50 mM TEAB, 0.1% SDS and 5 mM TCEP. 1:40 LysC enzyme was added and samples were incubated overnight at room temperature. Trypsin digestion was performed at a 1:70 enzyme-to-protein ratio for an 8-hour incubation at 37 °C.

### Proteoform identification

Quantitative reporter ion signals for PSMs were summarized to peptides by summation. Reporter ion signals of all individual temperatures were normalized using variance stabilizing normalization and converted to fold changes relative to the first temperature. Next, to assign similarly melting peptides found to map to a certain gene symbol, a graph for each gene symbol was created connecting all peptides (vertices) with weights (edges) corresponding to their similarity in melting profile. The similarity was computed using weighted euclidean distance, according to the formulas as described(*23*). Obtained graphs were then used for community detection using the Leiden algorithm. Only gene symbols for which at least ten peptides were identified and with at least two peptides per sample were used as input for graphs (a detected community had to be supported by at least three peptides to be accepted to ensure that outlier peptides did not affect robust proteoform identification). Peptides mapping to gene symbols for which these criteria were not fulfilled were grouped to single proteoforms, and peptides mapping to gene symbols that were included in the community detection were assigned to proteoform groups if the modularity of the detected communities was higher than 1 × 10−13 and the peptide ambiguity ratio was lower than 0.5 (for peptides mapping to multiple genes, it is calculated as the number of ambiguous peptides divided by the sum of the number of gene-specific and ambiguous peptides). Modularity was computed using the function modularity() of the igraph R package. Through the assignment of peptides to communities, functional proteoform groups for each gene symbol were created. Summarization on proteoform group level was performed by summation of non-normalized raw peptide data assigned to individual communities. Obtained proteoform signal intensities were then normalized per temperature using variance stabilizing normalization, and relative fold changes to the lowest measured temperature were formed.

Differential melting curve analysis was performed using NPARC, as previously published and described in the context of this analysis(*23*, *38*).

### Over-representation and network topology analysis

Over-Representation Analysis (ORA) and Network Topology-based Analysis (NTA) were performed using the R package WebGestaltR, version 0.4.6. The NTA was executed utilizing the WebGestaltR function, and Biological General Repository for Interaction Datasets (Biogrid) analysis was obtained utilizing the Homo Sapiens “network_PPI_BIOGRID” enrichment database, as included in the package. Genes symbols with an adjusted p-value (pAdj) below 0.05 were used as the genes of interest, with pAdj obtained from NPARC analysis of all melt curves and including only results with 8 full melting curves. The resultant network is formulated and prioritized with the “Network_Retrieval_Prioritization” method. Data was output for the top 100 results. Protein complex enrichment was performed using ORA with the Comprehensive Resource of Mammalian Protein Complexes (CORUM) database, also retrieved in the R package as the Homo Sapiens “network_CORUM” enrichment database. The full list of unique gene symbols that had 8 full melting curves were used as reference genes, and gene symbols with significant thermal melting changes as identified with NPARC were used as the hits input. Significance was determined using the false discovery rate (FDR) method Benjamini-Hochberg, “BH”, with a threshold of 0.05. A minimum number of 3 gene symbols in each CORUM complex category was required.

## Code availability

All code used to perform the computational analyses described and to reproduce the figures will be provided upon request and publicly available at time of peer reviewed publication.

## Contributions

R.J. designed the study, coordinated the study and provided the MS platform and funding. R.J., I.L., and E.K., developed the methodology and performed mass spectrometry experiments with help from A.A. and J.E.. E.K., A.A., and M.T. performed the TPP experiments. E.K. and J.E. performed the pull-down experiments. I.L. developed data analysis strategies and analyzed the data. I.L. drafted the manuscript with input from R.J. and which was reviewed and edited by all authors. R.J. supervised the study.

## Acknowledgements

This study was supported by grants from the Swedish Childhood Cancer Foundation (R.J., TJ2016-0035, PR2016-0019, PR2019-0025 and PR2022-0009); the Swedish Research Council (R.J., 2017-01653); the Felix Mindus Contribution to Leukemia Research (R.J.); the Dr. Åke Olsson Foundation for Hematological Research (2021-00130); and Cancer Society Stockholm and the King Gustaf V Jubilee Fund (R.J., 204092 and 221132); R.J. acknowledge the Karolinska Institutet and Science for Life Laboratory. The authors would also like to thank Luay Aswad, Xuekang Qi, Henri Colyn Bwanika and Janne Lehtiö for valuable input.

## References

1. R. Aebersold, J. N. Agar, I. J. Amster, M. S. Baker, C. R. Bertozzi, E. S. Boja, C. E. Costello, B. F. Cravatt, C. Fenselau, B. A. Garcia, Y. Ge, J. Gunawardena, R. C. Hendrickson, P. J. Hergenrother, C. G. Huber, A. R. Ivanov, O. N. Jensen, M. C. Jewett, N. L. Kelleher, L. L. Kiessling, N. J. Krogan, M. R. Larsen, J. A. Loo, R. R. Ogorzalek Loo, E. Lundberg, M. J. MacCoss, P. Mallick, V. K. Mootha, M. Mrksich, T. W. Muir, S. M. Patrie, J. J. Pesavento, S. J. Pitteri, H. Rodriguez, A. Saghatelian, W. Sandoval, H. Schlüter, S. Sechi, S. A. Slavoff, L. M. Smith, M. P. Snyder, P. M. Thomas, M. Uhlén, J. E. Van Eyk, M. Vidal, D. R. Walt, F. M. White, E. R. Williams, T. Wohlschlager, V. H. Wysocki, N. A. Yates, N. L. Young, B. Zhang, How many human proteoforms are there? Nat. Chem. Biol. 14, 206–214 (2018).

2. C. Liu, R. Karam, Y. Zhou, F. Su, Y. Ji, G. Li, G. Xu, L. Lu, C. Wang, M. Song, J. Zhu, Y. Wang, Y. Zhao, W. C. Foo, M. Zuo, M. A. Valasek, M. Javle, M. F. Wilkinson, Y. Lu, The UPF1 RNA surveillance gene is commonly mutated in pancreatic adenosquamous carcinoma. Nat. Med. 20, 596–598 (2014).

3. K. Yoshida, M. Sanada, Y. Shiraishi, D. Nowak, Y. Nagata, R. Yamamoto, Y. Sato, A. Sato-Otsubo, A. Kon, M. Nagasaki, G. Chalkidis, Y. Suzuki, M. Otsu, N. Obara, M. Sakata-Yanagimoto, K. Ishiyama, H. Mori, F. Nolte, W.-K. Hofmann, S. Miyawaki, S. Sugano, C. Haferlach, H. Phillip Koeffler, L.-Y. Shih, T. Haferlach, S. Chiba, H. Nakauchi, S. Miyano, S. Ogawa, Frequent Pathway Mutations of Splicing Machinery in Myelodysplasia. Blood. 118 (2011), pp. 458–458.

4. D. Ruggero, Translational Control in Cancer Etiology. Cold Spring Harbor Perspectives in Biology. 4 (2012), pp. a015891–a015891.

5. F. M. Ferguson, N. S. Gray, Kinase inhibitors: the road ahead. Nat. Rev. Drug Discov. 17, 353–377 (2018).

6. R. C. Centore, G. J. Sandoval, L. M. M. Soares, C. Kadoch, H. M. Chan, Mammalian SWI/SNF Chromatin Remodeling Complexes: Emerging Mechanisms and Therapeutic Strategies. Trends in Genetics. 36 (2020), pp. 936–950.

7. E. Jacinto, R. Loewith, A. Schmidt, S. Lin, M. A. Rüegg, A. Hall, M. N. Hall, Mammalian TOR complex 2 controls the actin cytoskeleton and is rapamycin insensitive. Nat. Cell Biol. 6, 1122–1128 (2004).

8. L. C. Kim, R. S. Cook, J. Chen, mTORC1 and mTORC2 in cancer and the tumor microenvironment. Oncogene. 36, 2191–2201 (2017).

9. J. C. Boulos, M. R. Yousof Idres, T. Efferth, Investigation of cancer drug resistance mechanisms by phosphoproteomics. Pharmacol. Res. 160, 105091 (2020).

10. K. Kitamura, K. Nimura, Regulation of RNA Splicing: Aberrant Splicing Regulation and Therapeutic Targets in Cancer. Cells. 10 (2021), doi:10.3390/cells10040923.

11. M. Cerezo, C. Robert, L. Liu, S. Shen, The Role of mRNA Translational Control in Tumor Immune Escape and Immunotherapy Resistance. Cancer Research. 81 (2021), pp. 5596– 5604.

12. Y. Li, N. Sun, Z. Lu, S. Sun, J. Huang, Z. Chen, J. He, Prognostic alternative mRNA splicing signature in non-small cell lung cancer. Cancer Letters. 393 (2017), pp. 40–51.

13. A. Lorentzian, A. Uzozie, P. F. Lange, Origins and clinical relevance of proteoforms in pediatric malignancies. Expert Review of Proteomics. 16 (2019), pp. 185–200.

14. J. E. Melnyk, V. Steri, H. G. Nguyen, Y. C. Hwang, J. D. Gordan, B. Hann, F. Y. Feng, K. M. Shokat, Targeting a splicing-mediated drug resistance mechanism in prostate cancer by inhibiting transcriptional regulation by PKCβ1. Oncogene. 41, 1536–1549 (2022).

15. P. I. Poulikakos, Y. Persaud, M. Janakiraman, X. Kong, C. Ng, G. Moriceau, H. Shi, M. Atefi, B. Titz, M. T. Gabay, M. Salton, K. B. Dahlman, M. Tadi, J. A. Wargo, K. T. Flaherty, M. C. Kelley, T. Misteli, P. B. Chapman, J. A. Sosman, T. G. Graeber, A. Ribas, R. S. Lo, N. Rosen, D. B. Solit, RAF inhibitor resistance is mediated by dimerization of aberrantly spliced BRAF(V600E). Nature. 480 (2011), pp. 387–390.

16. D. B. Johnson, A. M. Menzies, L. Zimmer, Z. Eroglu, F. Ye, S. Zhao, H. Rizos, A. Sucker, R. A. Scolyer, R. Gutzmer, H. Gogas, R. F. Kefford, J. F. Thompson, J. C. Becker, C. Berking, F. Egberts, C. Loquai, S. M. Goldinger, G. M. Pupo, W. Hugo, X. Kong, L. A. Garraway, J. A. Sosman, A. Ribas, R. S. Lo, G. V. Long, D. Schadendorf, Acquired BRAF inhibitor resistance: A multicenter meta-analysis of the spectrum and frequencies, clinical behaviour, and phenotypic associations of resistance mechanisms. European Journal of Cancer. 51 (2015), pp. 2792–2799.

17. Z. Blanchard, B. T. Paul, B. Craft, W. M. ElShamy, BRCA1-IRIS inactivation overcomes paclitaxel resistance in triple negative breast cancers. Breast Cancer Res. 17, 5 (2015).

18. S. Klaeger, S. Heinzlmeir, M. Wilhelm, H. Polzer, B. Vick, P.-A. Koenig, M. Reinecke, B. Ruprecht, S. Petzoldt, C. Meng, J. Zecha, K. Reiter, H. Qiao, D. Helm, H. Koch, M. Schoof, G. Canevari, E. Casale, S. R. Depaolini, A. Feuchtinger, Z. Wu, T. Schmidt, L. Rueckert, W. Becker, J. Huenges, A.-K. Garz, B.-O. Gohlke, D. P. Zolg, G. Kayser, T. Vooder, R. Preissner, H. Hahne, N. Tõnisson, K. Kramer, K. Götze, F. Bassermann, J. Schlegl, H.-C. Ehrlich, S. Aiche, A. Walch, P. A. Greif, S. Schneider, E. R. Felder, J. Ruland, G. Médard, I. Jeremias, K. Spiekermann, B. Kuster, The target landscape of clinical kinase drugs. Science. 358 (2017), doi:10.1126/science.aan4368.

19. M. M. Savitski, F. B. M. Reinhard, H. Franken, T. Werner, M. F. Savitski, D. Eberhard, D. M. Molina, R. Jafari, R. B. Dovega, S. Klaeger, B. Kuster, P. Nordlund, M. Bantscheff, G. Drewes, Tracking cancer drugs in living cells by thermal profiling of the proteome. Science. 346 (2014), doi:10.1126/science.1255784.

20. J. C. Tran, L. Zamdborg, D. R. Ahlf, J. E. Lee, A. D. Catherman, K. R. Durbin, J. D. Tipton, A. Vellaichamy, J. F. Kellie, M. Li, C. Wu, S. M. M. Sweet, B. P. Early, N. Siuti, R. D. LeDuc, P. D. Compton, P. M. Thomas, N. L. Kelleher, Mapping intact protein isoforms in discovery mode using top-down proteomics. Nature. 480, 254–258 (2011).

21. L. M. Smith, N. L. Kelleher, Proteoforms as the next proteomics currency. Science. 359 (2018), pp. 1106–1107.

22. R. Aebersold, M. Mann, Mass-spectrometric exploration of proteome structure and function. Nature. 537, 347–355 (2016).

23. N. Kurzawa, I. R. Leo, M. Stahl, E. Kunold, I. Becher, A. Audrey, G. Mermelekas, W. Huber, A. Mateus, M. M. Savitski, R. Jafari, Deep thermal profiling for detection of functional proteoform groups. Nat. Chem. Biol. (2023), doi:10.1038/s41589-023-01284-8.

24. D. Martinez Molina, R. Jafari, M. Ignatushchenko, T. Seki, E. A. Larsson, C. Dan, L. Sreekumar, Y. Cao, P. Nordlund, Monitoring drug target engagement in cells and tissues using the cellular thermal shift assay. Science. 341, 84–87 (2013).

25. J. A. Burger, A. Tedeschi, P. M. Barr, T. Robak, C. Owen, P. Ghia, O. Bairey, P. Hillmen, N. L. Bartlett, J. Li, D. Simpson, S. Grosicki, S. Devereux, H. McCarthy, S. Coutre, H. Quach, G. Gaidano, Z. Maslyak, D. A. Stevens, A. Janssens, F. Offner, J. Mayer, M. O’Dwyer, A. Hellmann, A. Schuh, T. Siddiqi, A. Polliack, C. S. Tam, D. Suri, M. Cheng, F. Clow, L. Styles, D. F. James, T. J. Kipps, RESONATE-2 Investigators, Ibrutinib as Initial Therapy for Patients with Chronic Lymphocytic Leukemia. N. Engl. J. Med. 373, 2425–2437 (2015).

26. M. L. Wang, S. Rule, P. Martin, A. Goy, R. Auer, B. S. Kahl, W. Jurczak, R. H. Advani, J. E. Romaguera, M. E. Williams, J. C. Barrientos, E. Chmielowska, J. Radford, S. Stilgenbauer, M. Dreyling, W. W. Jedrzejczak, P. Johnson, S. E. Spurgeon, L. Li, L. Zhang, K. Newberry, Z. Ou, N. Cheng, B. Fang, J. McGreivy, F. Clow, J. J. Buggy, B. Y. Chang, D. M. Beaupre, L. A. Kunkel, K. A. Blum, Targeting BTK with ibrutinib in relapsed or refractory mantle-cell lymphoma. N. Engl. J. Med. 369, 507–516 (2013).

27. S. P. Treon, C. K. Tripsas, K. Meid, D. Warren, G. Varma, R. Green, K. V. Argyropoulos, G. Yang, Y. Cao, L. Xu, C. J. Patterson, S. Rodig, J. L. Zehnder, J. C. Aster, N. L. Harris, S. Kanan, I. Ghobrial, J. J. Castillo, J. P. Laubach, Z. R. Hunter, Z. Salman, J. Li, M. Cheng, F. Clow, T. Graef, M. L. Palomba, R. H. Advani, Ibrutinib in previously treated Waldenström’s macroglobulinemia. N. Engl. J. Med. 372, 1430–1440 (2015).

28. J. A. Woyach, A. S. Ruppert, D. Guinn, A. Lehman, J. S. Blachly, A. Lozanski, N. A. Heerema, W. Zhao, J. Coleman, D. Jones, L. Abruzzo, A. Gordon, R. Mantel, L. L. Smith, S. McWhorter, M. Davis, T.-J. Doong, F. Ny, M. Lucas, W. Chase, J. A. Jones, J. M. Flynn, K. Maddocks, K. Rogers, S. Jaglowski, L. A. Andritsos, F. T. Awan, K. A. Blum, M. R. Grever, G. Lozanski, A. J. Johnson, J. C. Byrd, BTKC481S-Mediated Resistance to Ibrutinib in Chronic Lymphocytic Leukemia. J. Clin. Oncol. 35, 1437–1443 (2017).

29. J. A. Woyach, R. R. Furman, T.-M. Liu, H. G. Ozer, M. Zapatka, A. S. Ruppert, L. Xue, D. H.-H. Li, S. M. Steggerda, M. Versele, S. S. Dave, J. Zhang, A. S. Yilmaz, S. M. Jaglowski, K. A. Blum, A. Lozanski, G. Lozanski, D. F. James, J. C. Barrientos, P. Lichter, S. Stilgenbauer, J. J. Buggy, B. Y. Chang, A. J. Johnson, J. C. Byrd, Resistance mechanisms for the Bruton’s tyrosine kinase inhibitor ibrutinib. N. Engl. J. Med. 370, 2286–2294 (2014).

30. A. Quinquenel, L.-M. Fornecker, R. Letestu, L. Ysebaert, C. Fleury, G. Lazarian, M.-S. Dilhuydy, D. Nollet, R. Guieze, P. Feugier, D. Roos-Weil, L. Willems, A.-S. Michallet, A. Delmer, K. Hormigos, V. Levy, F. Cymbalista, F. Baran-Marszak, French Innovative Leukemia Organization (FILO) CLL Group, Prevalence of BTK and PLCG2 mutations in a real-life CLL cohort still on ibrutinib after 3 years: a FILO group study. Blood. 134, 641–644 (2019).

31. K. J. Maddocks, A. S. Ruppert, G. Lozanski, N. A. Heerema, W. Zhao, L. Abruzzo, A. Lozanski, M. Davis, A. Gordon, L. L. Smith, R. Mantel, J. A. Jones, J. M. Flynn, S. M. Jaglowski, L. A. Andritsos, F. Awan, K. A. Blum, M. R. Grever, A. J. Johnson, J.C. Byrd, J. A. Woyach, Etiology of Ibrutinib Therapy Discontinuation and Outcomes in Patients With Chronic Lymphocytic Leukemia. JAMA Oncol. 1, 80–87 (2015).

32. J. A. Woyach, How I manage ibrutinib-refractory chronic lymphocytic leukemia. Blood. 129, 1270–1274 (2017).

33. B. R. Lanning, L. R. Whitby, M. M. Dix, J. Douhan, A. M. Gilbert, E. C. Hett, T. O. Johnson, C. Joslyn, J. C. Kath, S. Niessen, L. R. Roberts, M. E. Schnute, C. Wang, J. J. Hulce, B. Wei, L. O. Whiteley, M. M. Hayward, B. F. Cravatt, A road map to evaluate the proteome-wide selectivity of covalent kinase inhibitors. Nat. Chem. Biol. 10, 760–767 (2014).

34. J. Chen, T. Kinoshita, J. Sukbuntherng, B. Y. Chang, L. Elias, Ibrutinib Inhibits ERBB Receptor Tyrosine Kinases and HER2-Amplified Breast Cancer Cell Growth. Mol. Cancer Ther. 15, 2835–2844 (2016).

35. D. Miklos, C. S. Cutler, M. Arora, E. K. Waller, M. Jagasia, I. Pusic, M. E. Flowers, A. C. Logan, R. Nakamura, B. R. Blazar, Y. Li, S. Chang, I. Lal, J. Dubovsky, D. F. James, L. Styles, S. Jaglowski, Ibrutinib for chronic graft-versus-host disease after failure of prior therapy. Blood. 130, 2243–2250 (2017).

36. M. Ghandi, F. W. Huang, J. Jané-Valbuena, G. V. Kryukov, C. C. Lo, E. R. McDonald 3rd, J. Barretina, E. T. Gelfand, C. M. Bielski, H. Li, K. Hu, A. Y. Andreev-Drakhlin, J. Kim, J. M. Hess, B. J. Haas, F. Aguet, B. A. Weir, M. V. Rothberg, B. R. Paolella, M. S. Lawrence, R. Akbani, Y. Lu, H. L. Tiv, P. C. Gokhale, A. de Weck, A. A. Mansour, C. Oh, J. Shih, K. Hadi, Y. Rosen, J. Bistline, K. Venkatesan, A. Reddy, D. Sonkin, M. Liu, J. Lehar, J. M. Korn, D. A. Porter, M. D. Jones, J. Golji, G. Caponigro, J. E. Taylor, C. M. Dunning, A. L. Creech, A. C. Warren, J. M. McFarland, M. Zamanighomi, A. Kauffmann, N. Stransky, M. Imielinski, Y. E. Maruvka, A. D. Cherniack, A. Tsherniak, F. Vazquez, J. D. Jaffe, A. A. Lane, D. M. Weinstock, C. M. Johannessen, M. P. Morrissey, F. Stegmeier, R. Schlegel, W. C. Hahn, G. Getz, G. B. Mills, J. S. Boehm, T. R. Golub, L. A. Garraway, W. R. Sellers, Next-generation characterization of the Cancer Cell Line Encyclopedia. Nature. 569, 503–508 (2019).

37. R. M. M. Branca, L. M. Orre, H. J. Johansson, V. Granholm, M. Huss, Å. Pérez-Bercoff, J. Forshed, L. Käll, J. Lehtiö, HiRIEF LC-MS enables deep proteome coverage and unbiased proteogenomics. Nat. Methods. 11, 59–62 (2014).

38. D. Childs, K. Bach, H. Franken, S. Anders, N. Kurzawa, M. Bantscheff, M. M. Savitski, W. Huber, Nonparametric Analysis of Thermal Proteome Profiles Reveals Novel Drug-binding Proteins. Mol. Cell. Proteomics. 18, 2506–2515 (2019).

39. F. Meissner, J. Geddes-McAlister, M. Mann, M. Bantscheff, The emerging role of mass spectrometry-based proteomics in drug discovery. Nat. Rev. Drug Discov. 21, 637–654 (2022).

40. G. Tsitsiridis, R. Steinkamp, M. Giurgiu, B. Brauner, G. Fobo, G. Frishman, C. Montrone, A. Ruepp, CORUM: the comprehensive resource of mammalian protein complexes-2022. Nucleic Acids Res. 51, D539–D545 (2023).

41. H. J. kleine Balderhaar, J. Lachmann, E. Yavavli, C. Bröcker, A. Lürick, C. Ungermann, The CORVET complex promotes tethering and fusion of Rab5/Vps21-positive membranes. Proceedings of the National Academy of Sciences. 110, 3823–3828 (2013).

42. G. Segala, M. A. Bennesch, N. M. Ghahhari, D. P. Pandey, P. C. Echeverria, F. Karch, R. K. Maeda, D. Picard, Vps11 and Vps18 of Vps-C membrane traffic complexes are E3 ubiquitin ligases and fine-tune signalling. Nat. Commun. 10, 1833 (2019).

43. S. Kumar, Z. Zeng, A. Bagati, R. E. Tay, L. A. Sanz, S. R. Hartono, Y. Ito, F. Abderazzaq, E. Hatchi, P. Jiang, A. N. R. Cartwright, O. Olawoyin, N. D. Mathewson, J. W. Pyrdol, M. Z. Li, J. G. Doench, M. A. Booker, M. Y. Tolstorukov, S. J. Elledge, F. Chédin, X. S. Liu, K. W. Wucherpfennig, CARM1 Inhibition Enables Immunotherapy of Resistant Tumors by Dual Action on Tumor Cells and T Cells. Cancer Discov. 11, 2050–2071 (2021).

44. M. Santos, J. W. Hwang, M. T. Bedford, CARM1 arginine methyltransferase as a therapeutic target for cancer. J. Biol. Chem. 299, 105124 (2023).

45. I. G. Solman, L. K. Blum, H. Y. Hoh, T. J. Kipps, J. A. Burger, J. C. Barrientos, S. O’Brien, S. P. Mulligan, N. E. Kay, P. Hillmen, J. C. Byrd, I. D. Lal, J. P. Dean, A. Mongan, Ibrutinib restores immune cell numbers and function in first-line and relapsed/refractory chronic lymphocytic leukemia. Leuk. Res. 97, 106432 (2020).

46. M. Long, K. Beckwith, P. Do, B. L. Mundy, A. Gordon, A. M. Lehman, K. J. Maddocks, C. Cheney, J. A. Jones, J. M. Flynn, L. A. Andritsos, F. Awan, J. A. Fraietta, C. H. June, M. V. Maus, J. A. Woyach, M. A. Caligiuri, A. J. Johnson, N. Muthusamy, J. C. Byrd, Ibrutinib treatment improves T cell number and function in CLL patients. J. Clin. Invest. 127, 3052– 3064 (2017).

47. J. Pan, R. M. Meyers, B. C. Michel, N. Mashtalir, A. E. Sizemore, J. N. Wells, S. H. Cassel, F. Vazquez, B. A. Weir, W. C. Hahn, J. A. Marsh, A. Tsherniak, C. Kadoch, Interrogation of Mammalian Protein Complex Structure, Function, and Membership Using Genome-Scale Fitness Screens. Cell Syst. 6, 555–568.e7 (2018).

48. R. D. Smith, V. V. Lupashin, Role of the conserved oligomeric Golgi (COG) complex in protein glycosylation. Carbohydr. Res. 343, 2024–2031 (2008).

49. M. Ishii, V. V. Lupashin, A. Nakano, Detailed Analysis of the Interaction of Yeast COG Complex. Cell Struct. Funct. 43, 119–127 (2018).

50. S. Drennan, G. Chiodin, A. D’Avola, I. Tracy, P. W. Johnson, L. Trentin, A. J. Steele, G. Packham, F. K. Stevenson, F. Forconi, Ibrutinib Therapy Releases Leukemic Surface IgM from Antigen Drive in Chronic Lymphocytic Leukemia Patients. Clin. Cancer Res. 25, 2503– 2512 (2019).

51. B. Simonetti, P. J. Cullen, Actin-dependent endosomal receptor recycling. Curr. Opin. Cell Biol. 56, 22–33 (2019).

52. C. M. Buckley, N. Gopaldass, C. Bosmani, S. A. Johnston, T. Soldati, R. H. Insall, J. S. King, WASH drives early recycling from macropinosomes and phagosomes to maintain surface phagocytic receptors. Proc. Natl. Acad. Sci. U. S. A. 113, E5906–E5915 (2016).

53. S.-S. Chen, B. Y. Chang, S. Chang, T. Tong, S. Ham, B. Sherry, J. A. Burger, K. R. Rai, N. Chiorazzi, BTK inhibition results in impaired CXCR4 chemokine receptor surface expression, signaling and function in chronic lymphocytic leukemia. Leukemia. 30, 833–843 (2016).

54. A. Bercusson, T. Colley, A. Shah, A. Warris, D. Armstrong-James, Ibrutinib blocks Btk-dependent NF-ĸB and NFAT responses in human macrophages during Aspergillus fumigatus phagocytosis. Blood. 132, 1985–1988 (2018).

55. D. Blez, M. Blaize, C. Soussain, A. Boissonnas, A. Meghraoui-Kheddar, N. Menezes, A. Portalier, C. Combadière, V. Leblond, D. Ghez, A. Fekkar, Ibrutinib induces multiple functional defects in the neutrophil response against Aspergillus fumigatus. Haematologica. 105, 478–489 (2020).

56. Y. Schurr, L. Reil, M. Spindler, B. Nieswandt, L. M. Machesky, M. Bender, The WASH-complex subunit Strumpellin regulates integrin αIIbβ3 trafficking in murine platelets. Sci. Rep. 13, 9526 (2023).

57. G. Dobie, F. A. Kuriri, M. M. A. Omar, F. Alanazi, A. M. Gazwani, C. P. S. Tang, D. M.-Y. Sze, S. M. Handunnetti, C. Tam, D. E. Jackson, Ibrutinib, but not zanubrutinib, induces platelet receptor shedding of GPIb-IX-V complex and integrin αIIbβ3 in mice and humans. Blood Adv. 3, 4298–4311 (2019).

58. E. L. Huttlin, R. J. Bruckner, J. Navarrete-Perea, J. R. Cannon, K. Baltier, F. Gebreab, M. P. Gygi, A. Thornock, G. Zarraga, S. Tam, J. Szpyt, B. M. Gassaway, A. Panov, H. Parzen, S. Fu, A. Golbazi, E. Maenpaa, K. Stricker, S. Guha Thakurta, T. Zhang, R. Rad, J. Pan, D. P. Nusinow, J. A. Paulo, D. K. Schweppe, L. P. Vaites, J. W. Harper, S. P. Gygi, Dual proteome-scale networks reveal cell-specific remodeling of the human interactome. Cell. 184, 3022–3040.e28 (2021).

59. Y. Liao, J. Wang, E. J. Jaehnig, Z. Shi, B. Zhang, WebGestalt 2019: gene set analysis toolkit with revamped UIs and APIs. Nucleic Acids Res. 47, W199–W205 (2019).

60. S. A. Misek, P. A. Newbury, E. Chekalin, S. Paithankar, A. I. Doseff, B. Chen, K. A. Gallo, R. R. Neubig, Ibrutinib Blocks YAP1 Activation and Reverses BRAF Inhibitor Resistance in Melanoma Cells. Mol. Pharmacol. 101, 1–12 (2022).

61. N. P. D. Liau, T. J. Wendorff, J. G. Quinn, M. Steffek, W. Phung, P. Liu, J. Tang, F. J. Irudayanathan, S. Izadi, A. S. Shaw, S. Malek, S. G. Hymowitz, J. Sudhamsu, Negative regulation of RAF kinase activity by ATP is overcome by 14-3-3-induced dimerization. Nat. Struct. Mol. Biol. 27, 134–141 (2020).

62. F. A. Cook, S. J. Cook, Inhibition of RAF dimers: it takes two to tango. Biochem. Soc. Trans. 49, 237–251 (2021).

63. B. Mezquita, M. Reyes-Farias, M. Pons, Targeting the Src N-terminal regulatory element in cancer. Oncotarget. 14, 503–513 (2023).

64. T. Varughese, Y. Taur, N. Cohen, M. L. Palomba, S. K. Seo, T. M. Hohl, G. Redelman-Sidi, Serious Infections in Patients Receiving Ibrutinib for Treatment of Lymphoid Cancer. Clin. Infect. Dis. 67, 687–692 (2018).

65. A. Wang, X.-E. Yan, H. Wu, W. Wang, C. Hu, C. Chen, Z. Zhao, P. Zhao, X. Li, L. Wang, B. Wang, Z. Ye, J. Wang, C. Wang, W. Zhang, N. S. Gray, E. L. Weisberg, L. Chen, J. Liu, C.- H. Yun, Q. Liu, Ibrutinib targets mutant-EGFR kinase with a distinct binding conformation. Oncotarget. 7, 69760–69769 (2016).

66. R. G. Newcombe, Interval estimation for the difference between independent proportions: comparison of eleven methods. Stat. Med. 17, 873–890 (1998).

67. T. Paysan-Lafosse, M. Blum, S. Chuguransky, T. Grego, B. L. Pinto, G. A. Salazar, M. L. Bileschi, P. Bork, A. Bridge, L. Colwell, J. Gough, D. H. Haft, I. Letunić, A. Marchler-Bauer, H. Mi, D. A. Natale, C. A. Orengo, A. P. Pandurangan, C. Rivoire, C. J. A. Sigrist, I. Sillitoe, N. Thanki, P. D. Thomas, S. C. E. Tosatto, C. H. Wu, A. Bateman, InterPro in 2022. Nucleic Acids Res. 51, D418–D427 (2023).

